# Histone Acetylation Differentially Modulates CTCF-CTCF Loops and Intra-TAD Interactions

**DOI:** 10.1101/2025.07.29.667515

**Authors:** Rebecca G Smith, Yu Fu, Kathleen L Schiela, Madison Dautle, Ryan Williams, Hannah M Wilson, Chloe Azadegan, Johnathan R Whetstine, Job Dekker, Yu Liu

**Author notes:** Correspondence (Y.L.); (J.D.).

## Abstract

The cohesin complex structures the interphase genome by extruding loops and organizing topologically associating domains (TADs). While cohesin engages chromatin in context-dependent modes, the regulatory influence of chromatin state on these interactions remains unclear. Here, we show that histone hyperacetylation, induced by the histone deacetylase inhibitor trichostatin A (TSA), preferentially disrupts short-range interactions within TADs but spares CTCF-anchored loops, despite reduced cohesin occupancy at these sites. These findings point to two functionally distinct cohesin populations: a TSA-sensitive pool within TADs, likely representing extruding, non-topologically bound cohesin, and a TSA-resistant population at CTCF–CTCF anchors that maintains loops through topological entrapment. Using a semi–in vitro system with TEV-cleavable RAD21, we show that TSA-resistant cohesin at CTCF sites becomes TSA-sensitive after proteolytic cleavage that opens the cohesin ring, showing that it is the topological engagement with DNA that makes cohesin, and CTCF-CTCF loops, TSA-resistant. Notably, we also detect TSA-sensitive cohesin at CTCF sites, suggesting the presence of transient, non-encircling cohesin that either precedes conversion to the stable form or is halted by pre-existing encircling cohesin. Together, our results suggest that cohesin exists in distinct biochemical states: an extruding form found within TADs and at CTCF sites, that is sensitive to hyperacetylation, and a topologically bound form specifically at CTCF-CTCF loops that is insensitive. The former may allow dynamic changes in chromatin loops, while latter ensures robustness of CTCF-anchored loops in response to chromatin state changes.

## Introduction

The cohesin complex plays a critical role in the formation of chromatin loops and domains^1,2^ and thereby contributes to the three-dimensional (3D) organization of the genome. Cohesin is composed of three core subunits: SMC1, SMC3 and RAD21, which together form a ring structure^3–7^. This ring structure allows cohesin to mediate cohesion of sister chromatids during mitosis^7–9^ by topologically encircling two sister chromatids. During interphase, cohesin actively extrudes chromatin loops that bring distant genomic regions into proximity^10–12^. CTCF-bound sites block cohesin-mediated loop extrusion in an orientation-dependent manner, leading to stalling of cohesin complexes at these sites. These stalled cohesin complexes produce chromosome structures that are visible as dots of enriched interactions on Hi-C contact maps. In contrast, the actively extruding cohesin mediates transient interactions, not reproducibly positioned at specific positions. Over the population, the sum of these dynamic loops produce enriched interactions within topologically associating domains (TADs)^2,13–17^.

Recent studies have shown that the stalled cohesin at convergent CTCF sites entraps chromatin in a similar way as cohesive cohesin holds pairs of sister chromatids during mitosis, i.e., chromatin going through the ring^18,19^. Proteolytic cleavage of RAD21 leads to opening of the cohesin ring, cohesin dissociation from CTCF-bound sites, and loss of CTCF-CTCF loops. In contrast, actively extruding cohesin complexes that are not stalled at CTCF sites and that are positioned within TADs engage chromatin in a different way (chromatin doesn’t go through the cohesin ring). These extruding complexes “clamp” the chromatin template^20,21^, remain bound to chromatin after proteolytic cleavage of RAD21, and their loops remain in place. These complexes can be dissociated from DNA by high salt concentrations even without cleavage of RAD21 indicating they do not encircle the DNA. These findings indicate cohesin engages with chromatin in different ways depending on its location within the genome. However, the precise mechanisms by which cohesin interacts with chromatin in these varied contexts, and how these interactions are modulated by chromatin modifications, remain incompletely understood.

One such chromatin modification is histone acetylation, which is known to reduce nucleosome interactions and promote a more open chromatin conformation^22^. Histone hyperacetylation, e.g., induced in the presence of HDAC inhibitors like Trichostatin A (TSA), leads to global chromatin decompaction^23–25^. While the effects of histone hyperacetylation on chromatin accessibility and gene expression have been well documented, its impact on cohesin-mediated chromatin interactions is less clear.

In this study, we demonstrated that TSA-induced hyperacetylated histones have minimal impact on long-range chromatin interactions (>2 Mb) but reduce short-range (<2 Mb) interactions, due to loss of cohesin loops within TADs. While CTCF-CTCF loops are not affected by hyperacetylation when the cohesin ring is intact, they are lost to a greater extent after RAD21 cleavage. These findings suggest that histone acetylation lowers the affinity of the actively extruding clamped cohesin complexes within TADs, while cohesin complexes at the bases of CTCF-CTCF loops remain bound as long as they can encircle the two looping anchors. Most notably, our results suggest the presence of two distinct populations of cohesin at CTCF–CTCF sites: one that pseudotopologically encircles chromatin and is resistant to histone hyperacetylation (TSA-resistant), and another that does not encircle chromatin and is sensitive to histone hyperacetylation (TSA-sensitive), resembling the cohesin found within TADs. Together, these findings provide mechanistic insight into the context-dependent behavior of cohesin and underscore the differential impact of histone acetylation on 3D genome organization.

## Results

### Impact of TSA-induced histone hyperacetylation on genome compartmentalization

TSA is widely recognized for its ability to increase histone acetylation and promote chromatin decompaction^23,24,26^. While many studies employ overnight TSA treatments, such prolonged exposure often leads to extensive changes in the transcriptome, epigenome, and nuclear morphology, complicating the interpretation of results. To minimize these confounding effects, we treated cells with 500 nM TSA for 3 hours, as supported by recent studies^27,28^. Western blot analysis confirmed that histone acetylation levels peaked under these conditions (Extended Data Fig. 1a-b). Furthermore, cell cycle analysis indicated that this treatment did not perturb cell cycle progression (Extended Data Fig. 1c). Thus, we adopted the 3-hour 500 nM TSA treatment regimen for all subsequent experiments.

We next performed Hi-C analysis on HAP1-RAD21^TEV^ cells^18^ following TSA treatment. Hi-C contact maps revealed that TSA treatment increases the E1 eigenvector values at some loci (dark arrows, Fig. 1a-b; Extended Data Fig. 1d-e). The E1 eigenvector, derived from principal component analysis of the Hi-C interaction matrix, is a widely used metric to delineate A and B compartment positions^29^.

**Fig. 1.**
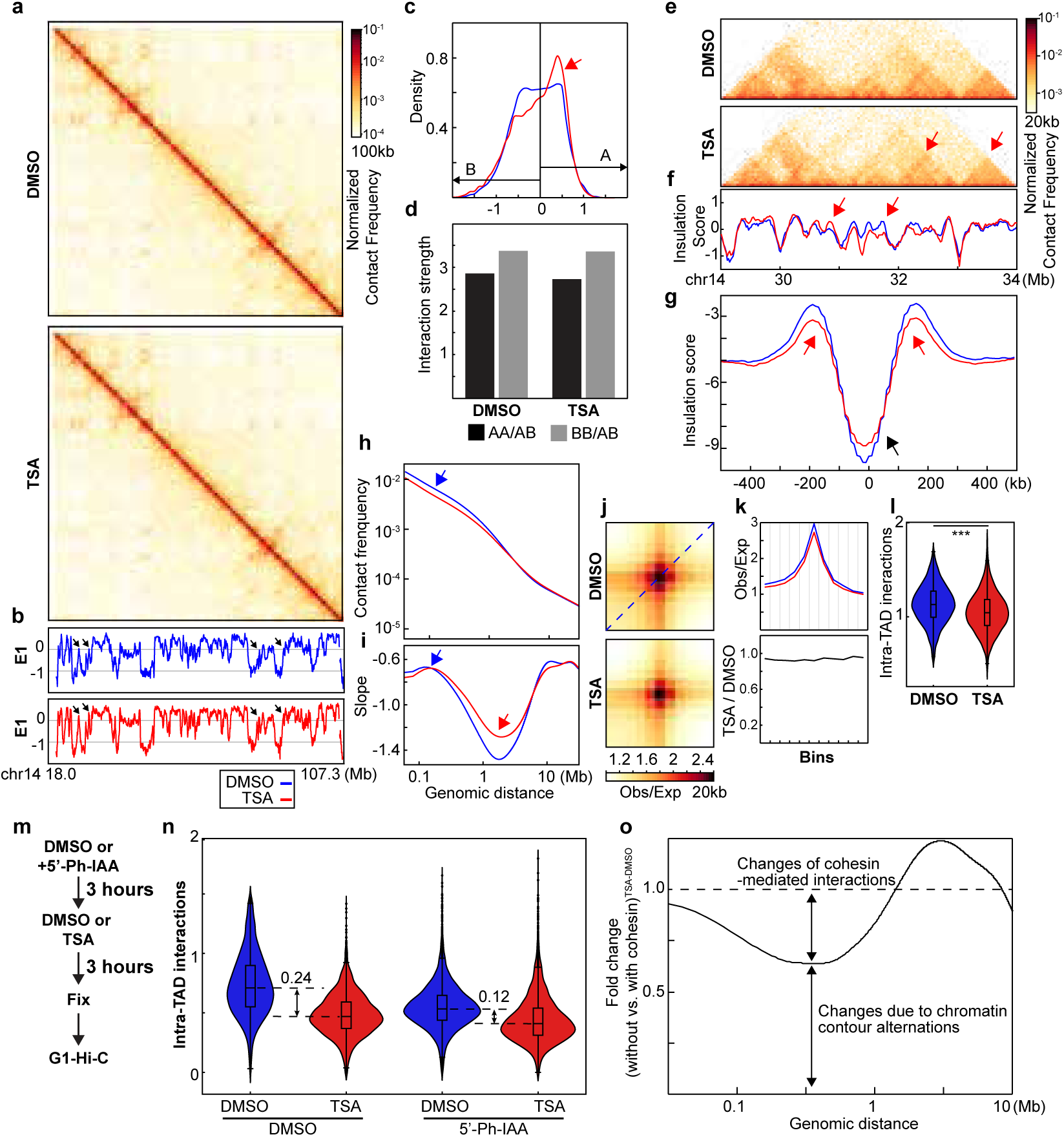
Histone hyperacetylation alters genome interactions. **a**, Hi-C interaction maps for HAP1 cells treated with/without TSA (top). Data are for the 18.0–107.3 Mb region of chromosome 14. **b**, Eigenvector E1 across the same region as in **a**. Dark arrows indicate loci with increased E1 values. **c**, Distribution of E1. The red arrow pointed the increased density of E1 bins (A compartment signals) after TSA treatment. **d,** Interaction strength of compartments. Dark and grey bars indicate the strength of the A-A and B-B interactions, respectively (see Methods). **e**, Hi-C interaction maps for HAP1 cells treated with/without TSA. Data for the 29–34 Mb region of chromosome 14 are shown. **f**, Insulation profiles for the same region as in **e**. **g**, Aggregate Hi-C data at TAD boundaries identified in the sample treated with DMSO. **h and i**, *P(s)* plots (**h**) and plots of their derivatives (**i**) for Hi-C data from cells treated with/without TSA. The arrow indicates the signature of cohesin loops. **j**, Aggregated Hi-C data at 8,334 loops identified in HAP1 cells according to^12^ (upper heatmap). **k**, The average Hi-C signals from the bottom-left corner to the top-right corner of the respective loop aggregation heatmaps (top), as illustrated by the blue dashed line in the top Hi-C panel in **j**. This line is defined as the loop-line. The blue and red loop-lines represent loop strength in DMSO and TSA treated samples, respectively. The dark line in the bottom plot indicates the ratio between TSA (red) and DMSO (blue) loop-lines. **l**, Average intra-TAD interactions across all TADs (see methods). Blue and red represent DMSO and TSA, respectively. Wilcoxon sum rank test, ***p<0.001. **m and n**, Calculation of TSA-induced reduction of cohesin-mediated intra-TAD interactions. RAD21 degron cells (HCT116-AID-RAD21) were firsted treated with 5’-Ph-IAA to eliminate cohesin 3 hours before DMSO or TSA treatment for subsequent three hours. The G1 cells were then sorted to perform Hi-C analysis (**m**) and short-range interactions within TADs, defined using cells with cohesin and treated with DMSO, were calculated and ploted (**n**). **o**, Fold change of TSA-induced *P(s)* alterations in the absence versus the presence of cohesin.

We next analyzed the distribution of E1 values and found that TSA-induced histone hyperacetylation increased the number of A compartment bins, consistent with observed elevation in E1 values and indicative of some changes in compartment patterns (Fig. 1c; Extended Data Fig. 1f). We calculated compartment strength, i.e., the strength of the preference of interactions between loci of the same compartment type, by ranking loci by their E1 value and plotting pairwise interactions normalized for their expected interaction frequency given their genomic distance. Compartment strength is then calculated as the ratio of A-A or B-B to A-B and B-A interactions (see Methods)^15^. Histone acetylation is known to promote chromatin decompaction, which could potentially influence compartment interactions. However, we observed minimal changes in compartmentalization strength in TSA-treated cells (Fig. 1d; Extended Data Fig. 1g). In summary, TSA treatment alters compartment patterns by increasing the number of A compartment bins but has minimal impact on overall compartment interaction strength.

### TSA-induced histone acetylation reduces local chromatin interactions

To investigate the local effects of TSA treatment on chromatin organization, we first analyzed Hi-C contact maps. TSA-treated cells displayed reduced interactions within chromatin domains (red arrows, Fig. 1e; Extended Data Fig. 1h). This can be quantified directly by plotting Hi-C interaction frequency (*P*) as a function of genomic distance (*s*) (Fig. 1g and Extended Data Fig. 1k). In TSA treated cells, interaction frequencies for loci separated by up to 2 Mb are reduced about 20%. At this length scale cohesin-mediated loops and TADs are dominant chromatin folding features. We therefore examined these features in more detail by calculating insulation profiles^30^ (Fig. 1f; Extended Data Fig. 1i) and observed a notable reduction in insulation scores at domain boundaries following TSA treatment. We next assessed TAD boundary insulation strength by aggregating interactions at and around boundaries. In DMSO-treated cells, the characteristic depletion of interactions across boundaries was evident, as shown by the depth of the valley in the insulation profile (blue line, Fig. 1g; Extended Data Fig. 1j), which reflects the strength of boundary insulation between adjacent TADs. In TSA-treated cells, the valley depth was reduced (red line, Fig. 1g; Extended Data Fig. 1j), indicating weakened TAD boundary insulation. This reduction likely stems from decreased interactions within flanking TADs, as reflected by the diminished shoulders of the valley (red arrows, Fig 1g and Extended Data Fig. 1j). In summary, TSA treatment reduces local chromatin interactions and weakens TADs.

### Histone hyperacetylation reduces cohesin-mediated loops

The shape of *P(s)* reflects key features of chromatin organization, including cohesin-mediated loops^31,32^. In DMSO-treated cells, *P(s)* displayed a characteristic shoulder where interactions decayed more slowly for loci separated by ∼100 kb (Fig. 1h; Extended Data Fig. 1k, blue arrow). The derivative of *P(s)* highlighted this feature as a local peak, corresponding to the average loop size, while the valley depth at ∼ 2Mb reflected loop densities (Fig. 1i and Extended Data Fig. 1l)^31,33^.

For TSA-treated cells, the derivative of *P(s)* showed two changes compared to controls: first, the local peak reflecting average loop size was slightly shifted to the right; and second, the depth of the valley at ∼2Mb was reduced (Fig. 1i; Extended Data Fig. 1l, red arrow). Together, this indicates a reduction of the number of cohesin-mediated loops, and coincident extension of their size. Cohesin mediates two major types of chromatin loops: loops within TADs generated by actively extruding cohesin and CTCF-CTCF loops stabilized by cohesin stalling at CTCF sites. The derivative plots represent signals from both types of cohesin-mediates loops^1,2,18,31,33^. The reduced peak depth at ∼2 Mb in TSA-treated cells suggests a loss of loops, but does not provide information about which loops are lost. To determine whether loops generated by cohesin that is actively extruding cohesin, or by cohesin stalled at pairs of CTCF sites, or both are affected, we next examined the impact on each loop type individually.

We first investigated the impact of TSA treatment on CTCF-CTCF looping interactions. In DMSO-treated cells, strong focal enrichment of interactions at loop anchors was readily observed, and this enrichment remained largely unchanged in TSA-treated cells (Fig. 1k and Extended Data Fig. 1n). To quantify loop strength, we plotted Hi-C interaction frequencies along the diagonal of loop aggregation heatmaps (Fig. 1k, blue dashed line, top heatmap). These ’loop-line’ plots revealed peaks corresponding to CTCF-CTCF loop interactions (Fig. 1k, blue line)^18^, with TSA-treated cells represented by the red loop-line. Ratio of the blue (DMSO) loop-line from the red (TSA) loop-line showed minimal differences in loop strength (Fig. 1k, bottom panel, dark line), indicating that TSA treatment has little impact on most CTCF-CTCF loop interactions. These results suggest that loops mediated by cohesin at pairs of CTCF sites are largely resistant to TSA-induced histone hyperacetylation.

We next quantified the changes in intra-TAD interactions by calculating the average interaction frequency within TADs (see Methods). We observed that TSA treatment significantly reduced intra-TAD interactions across replicates, as reflected by the observed decrease in TAD boundary insulation strength (Fig. 1l and Extended Data Fig. 1n).

Histone hyperacetylation reduces nucleosome interactions and decompact chromatin fiber^25,34^, increasing its contour length which may contribute to the reduction of intra-TAD interactions. To assess how much chromatin decompaction decreases intra-TAD interactions, we first depleted all cohesin using RAD21-degron cells^35^, followed by treatment with either DMSO or TSA. As cohesin loss eliminate TAD structures, TADs were defined using the DMSO-treated, cohesin-intact condition, and changes in short-range interactions were quantified within these regions (Fig. 1m). As expected, eliminating cohesin significantly reduces intra-TAD interactions (Fig. 1n and Extended Data Fig. 1o, two replicates, the first and third blue violin plots in each replicate). TSA treatment further decreases intra-TAD interactions in the chromosomes without cohesin (Fig. 1n and Extended Data Fig. 1o, the forth red violin plots in each replicate). We then compare the difference between TSA and DMSO treatments with or without cohesin degradation. The decrease in intra-TAD interactions in cells without cohesin account for ∼50% decrease of intra-TAD interactions in cells with cohesin. Therefore, our results showed that ∼50% of the reduction of intra-TAD interactions reduced by TSA is due to reduced cohesin-mediated interactions within TADs. Further, the level of intra-TAD interactions in TSA-treated cells is comparable whether cells express cohesin or not. This is consistent with the model that TSA treatment leads to both chromatin decompaction and loss of cohesin binding. In summary, these results suggest that cohesin within TADs is selectively sensitive to TSA-induced histone hyperacetylation.

We next assessed the reduction of cohesin-mediated short-range interactions across genomic distances using *P(s)* analysis. TSA-induced changes in *P(s)* were compared in the absence and presence of cohesin, and their ratio was plotted (see Methods; Fig. 1o and Extended Data Fig. 1p for two replicates. The corresponding *P(s)* plots are shown in Extended Data Fig. 1q–r). Interestingly, the reduction in contact frequency resulting from chromatin contour length changes varied with genomic distance, showing minimal impact around 300 kb and maximal impact near 2 Mb. Since most cohesin-mediated intra-TAD interactions occur within 500 kb, our results suggest that the reduction of cohesin-mediated interactions accounts for approximately 40% of the observed reduction in interaction frequency. Notably, the *P(s)* ratio (absence vs. presence of cohesin) exceeds 1 in the >2 Mb range, indicating that loss of short-range loops leads to less decay at larger distance.

Taken together, our findings demonstrate that TSA-induced histone hyperacetylation reduces both nucleosome and cohesin-mediated looping interactions, resulting in significantly weakened intra-TAD interactions while having minimal impact on CTCF-CTCF loops.

SMC3 acetylation, which is regulated by HDAC8^36^, could also be modulated by TSA, potentially affecting cohesin-mediated interactions. To test this, we examined both the levels of acetylated SMC3 and the chromatin affinities of SMC3 and acetylated SMC3 following TSA treatment. We observed no significant changes in acetylated SMC3 levels or its chromatin affinities (Extended Data Fig. 2a). These results suggest that the loss of cohesin-mediated loops induced by TSA treatment is not attributable to changes in SMC3 acetylation.

In HAP1-RAD21^TEV^ cells, RAD21 contains a tandem TEV motif in the unstructured region^18^, which could potentially influence cohesin-mediated loops and contribute to the changes observed following TSA treatment. To assess the impact of TSA treatment on looping interactions, we examined HAP1 wild-type cells and found that TSA treatment led to similar effects on both compartmentalization and cohesin loops. These results indicate that the inserted TEV sequence in HAP1-RAD21TEV cells does not account for the observed changes following TSA treatment (Extended Data Figs. 2 and 3).

### Impact of histone hyperacetylation on chromatin binding of cohesin and CTCF

Histone acetylation alters the charge properties of chromatin, it may thus alter cohesin and chromatin interactions and disrupt loop extrusion, leading to the loss of cohesin-mediated loops within TADs. We thus used an assay we developed previously that can examine how TSA treatment, likely histone hyperacetylation, affects interactions between cohesin and chromatin^18^ (Fig. 3a). First, we investigated whether TSA affects chromatin binding of cohesin using purified nuclei from DMSO- or TSA-treated cells. Western blotting confirmed that histone acetylation induced by TSA treatment was preserved in the purified nuclei (Fig. 2a and Extended Data Fig. 5a), indicating that chromatin states in purified nuclei are comparable to those in intact cells. We then added TEV protease to the purified nuclei and cleaved RAD21 under low or physiological salt concentrations and assessed the chromatin association of cohesin subunits. Surprisingly, western blotting indicated that TSA treatment had minimal effects on the chromatin retention of both CTCF and cohesin core subunits, including RAD21, SMC1, SMC3, SA1 and SA2, in all the various conditions (Fig. 2b and Extended Data Fig. 5a). These results suggest that, despite histone hyperacetylation reducing cohesin-mediated loops within TADs, the global chromatin binding of cohesin core components remains unchanged.

**Fig. 2.**
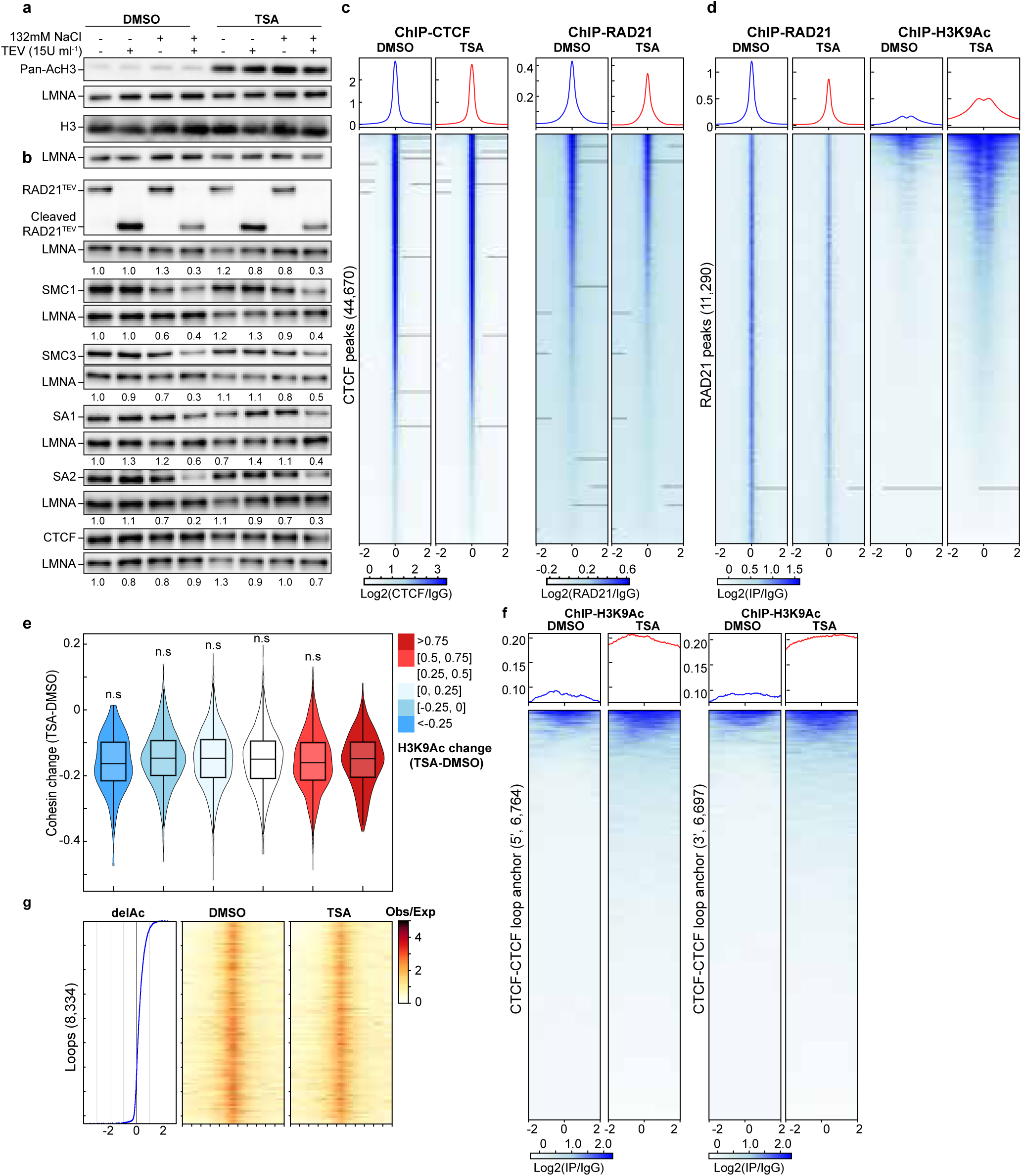
Impact of histone hyperacetylation on chromatin binding of CTCF and cohesin, and CTCF-CTCF loop anchors. **a**, The Western blots indicated levels of whole histone H3 and acetylated histones by a pan-acetylated histone antibody. **b**, Western blots of chromatin association of cohesin subunits in DMSO- or TSA-treated HAP1-RAD21^TEV^ nuclei in NB or NBS1 buffer (NB buffer + 132 mM NaCl) treated with/without TEV. LMNA was used for normalization (left). The levels of the cohesin subunits in each condition were normalized to their levels in DMOS-treated nuclei in NB buffer without TEV treatment and are indicated in under each LMNA blot. **c**, ChIP–seq signals of CTCF (left) and RAD21 (right) at CTCF binding sites. Average CTCF and RAD21 ChIP–seq signals in each condition for 44,670 CTCF binding sites identified from the CTCF ChIP data from cells treated with DMSO. Heatmaps of CTCF (left) and RAD21 (right) ChIP–seq signals for each condition at each CTCF binding site (bottom). **d**, ChIP–seq signals of RAD (left) and H3K9Ac (right) at RAD21 binding sites. Average RAD21 and H3K9Ac ChIP–seq signals in each condition for 11,290 RAD21 binding sites identified from the RAD21 ChIP data from cells treated with DMSO (top). Heatmaps of RAD21 (left) and H3K9Ac (right) ChIP– seq signals for each condition at each RAD21 binding site (bottom). **e**, Relationship between RAD21 binding changes and histone acetylation changes. Average of H3K9Ac ChIP-seq signal changes at RAD21 binding sites (+/-2kb) are binned and the changes RAD21 binding strength of each peak in each bin are plotted as a violin plot embed with a boxplot. **f**, ChIP–seq signals of H3K9Ac at 5’- and 3’-loop anchors. There are 6,764 5’- and 6,697 3’-loop unique anchors, which consist of 8,334 loops^12^. **g**, Relationship between histone acetylation changes and loop-line changes. Left ranking blue plot: combined changes of H3KAc ChIP-seq signals at both anchors of 8,334 loop sites^12^. Heatmap of ranked 8,334 loop-lines for each condition. n.s. not significant, Wilcoxon sum rank test.

**Fig. 3.**
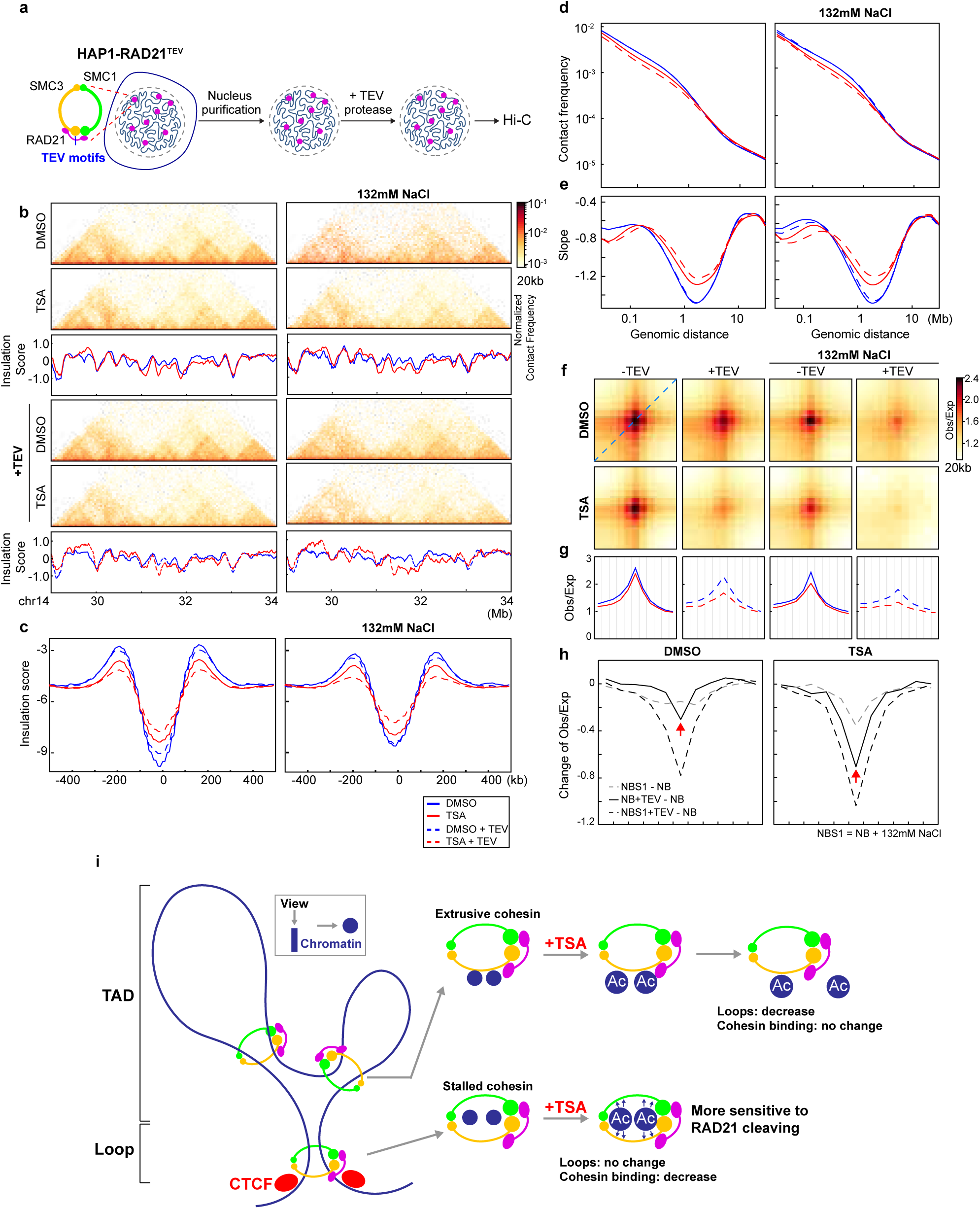
Histone hyperacetylation increases the sensitivity of cohesin at CTCF-CTCF loop anchors to RAD21 cleavage. **a**, A schemetic of semi-in vitro assay. Tandam TEV motifs were inserted to RAD21 using CRISPR to generate HAP1RAD21-TEV cells^18^. Nuclei were purified from HAP1RAD21- TEV cells and treated with TEV protease in different salt concentrations before Hi-C or Western blotting analysis. **b**, Examples of Hi-C maps obtained with DMSO- or TSA-treated HAP1-RAD21^TEV^ nuclei treated with or without TEV in NB (left) or NBS buffer (right). Insulation profiles (bottom) for the same region for each condition. **c**, Aggregate Hi-C data at TAD boundaries identified in DMSO- or TSA-treated nuclei treated with or without TEV in NB (left) or NBS1 (right) buffer. **d** and **e**, *P(s)* plots (**d**) and plots of their derivatives (**e**) for Hi-C data from DMSO- or TSA-treated nuclei with or without TEV treatment in NB (left) or NBS1 (right) buffers. The arrows indicate the signature of cohesin loops. **f**, Aggregate Hi-C data at loops identified in HAP1 cells (as in Fig. 1j; top). **g,** Loop-lines for each condition. In each panel, red and blue lines indicate DMSO or TSA treatments and dash lines indicate TEV protease treatment. **h**, The difference of loop-lines (all other conditions versus NB buffer), left and right panels are DMSO versus TSA treatment. **i,** A summary of distinct responses of cohesin-mediated loop interactions within TADs and at CTCF-CTCF sites. Cohesin-mediated looping interactions within TADs are reduced upon histone hyperacetylation, while cohesin loops at CTCF-CTCF binding sites are resistant to histone hyperacetylation. However, chromatin with hyperacetylated histones generates tension on cohesin ring structure (grey arrows), which is more sensitive to RAD21 cleavage. The dark blue dots indicate chromatin seen from the top.

We next performed ChIP-seq analysis to investigate the effects of histone hyperacetylation on CTCF and cohesin binding to chromatin (Fig. 2a and Extended Data Fig. 5a). Unlike Western-based methods that assess global chromatin binding, ChIP-seq specifically profiles enriched binding sites, particularly cohesin occupancy at CTCF sites.

While CTCF binding at CTCF sites was unaffected by TSA treatment, RAD21 ChIP signals at CTCF-bound sites showed some reduction. Intriguingly, this reduction occurred despite the relative stability of CTCF-mediated looping interactions under conditions of histone hyperacetylation. Further, RAD21 ChIP signals also showed a general decrease in cohesin chromatin association following histone acetylation (Fig. 2b and Extended Data Fig. 5b), indicating a general reduction of cohesin binding at specific sites.

We next assessed histone hyperacetylation. We focused on H3K9Ac ChIP signals, which are enriched in promoter and open chromatin regions^37^. TSA treatment led to a ∼50% increase in H3K9Ac peaks (Fig. 2b and Extended Data Fig. 5b, right panels). However, we did not detect a direct correlation between the increased H3K9Ac ChIP signals and the decrease in RAD21 ChIP signals (Fig. 2c and Extended Data Fig. 5c).

We then examined changes in H3K9Ac ChIP signals at CTCF-CTCF loop anchors. TSA treatment inreased H3K9Ac levels at both 5’- and 3’- loop anchors (Fig. 2f and Extended Data Fig. 4e), although the majority of anchors showed minimal changes. We next ranked loop anchors by degree of H3K9Ac increases and compared loop line signals between DMSO- and TSA-treated cells (Fig. 2e and Extended Data Fig. 4d). A modest reduction in loop line signals was observed upon TSA treatment, as reflected by the loop-line pileup analysis (Fig. 1k, 2f, Extended Data Fig. 1m, Extended Data Fig. 4f). However, this reduction did not scale with H3K9Ac levels. Together, these results confirm that CTCF-anchored loops are largely maintained despite histone hyperacetylation.

Taken together, our results indicated that cohesin binding showed minimal changes within TADs, while cohesin-mediated loops within TADs are reduced. This suggests that although cohesin can still associate with acetylated chromatin within TADs, its ability to tether two loci and mediate loop formation is reduced. In contrast, cohesin binding at CTCF-CTCF sites is reduced, while CTCF-CTCF looping interactions remain largely unchanged.

### Histone hyperacetylation increases the sensitivity of CTCF looping interactions to cohesin ring integrity

Others and we have showed that cohesin entrapped chromatin at CTCF-CTCF sites, this may contribute to stability of loop interactions at CTCF-CTCF sites after TSA treatment.

We thus sought to explore how histone hyperacetylation influences cohesin association with chromatin at CTCF-CTCF loop sites. The semi-in vitro nuclear system, established previously, enables controlled perturbations of cohesin interactions with chromatin^18^, making it ideal for measuring cohesin-chromatin interactions at CTCF-CTCF loop sites following TSA treatment (Fig. 3a). We purified nuclei from DMSO- or TSA-treated cells, then cleaved RAD21 under low or physiological salt conditions to assess the cohesin engagement with chromatin at these sites.

Hi-C maps revealed that TSA treatment did not cause significant alterations in compartment boundaries or compartment interaction strength, consistent with the Hi-C analysis in cells (Extended Data Fig. 7b-c and Extended Data Fig. 8a-b). We then examined local features of chromosome structure following TSA treatment. Hi-C maps showed that RAD21 cleavage under both salt buffers led to minimal changes in local chromatin structures (Fig. 3c and Extended Data Fig. 8c). As anticipated, TSA treatment induced notable changes in certain local chromatin features (Fig. 3c and Extended Data Fig. 8c). To quantify the loss of local features, we calculated insulation profiles (Fig. 3c and Extended Data Fig. 8c, blue versus red lines, representing DMSO and TSA treatments, respectively). Consistent with our Hi-C analysis in cells, we observed significant changes in some domains, including changes in domain boundary strength following TSA treatment.

We then assessed the insulation strength by aggregating interactions at and around TAD boundaries. Consistent with previous studies^18^, both salt treatment and RAD21 cleavage weakened the insulation strength, as indicated by the blue plots in both panels (Fig. 3d and Extended Data Fig. 8d). Interestingly, TSA treatment also weakened insulation strength at TAD boundaries, as evidenced by the red plots in both panels (Fig. 3d and Extended Data Fig. 8d). These synergistic effects suggest that the impact of TSA on genome folding is distinct from the effects of salt or RAD21 cleavage, highlighting a unique mode of action for TSA in chromatin organization.

We then analyzed how interaction frequency decayed with genomic distance. Interestingly, short-range interactions (<1 Mb) in TSA-treated nuclei were more sensitive to RAD21 cleavage (Fig. 3e and Extended Data Fig. 8e, dashed red lines), while long-range interactions (>1 Mb) showed minimal changes across TSA, RAD21 cleavage, and salt treatments. These differences are further highlighted in the derivative of *P(s)*, where cleaving RAD21 in TSA-treated nuclei resulted in more pronounced changes to cohesin density peaks, as evidenced by the difference between the solid and dashed red lines compared to the blue lines (Fig. 3f and Extended Data Fig. 8f). These results indicated that TSA treatment mostly reduces cohesin-mediated loops within TADs, consistent with our living cell results, while with TSA and cleavage, all loops are reduced including non-extruding loops at CTCF-CTCF sites.

In our previous studies, we observed that CTCF-CTCF loops are sensitive to RAD21 cleavage, whereas intra-TAD loops are resistant^18^. Based on this, we hypothesize that histone hyperacetylation changes the chromatin affinity of cohesin at CTCF-CTCF loop sites, increasing their sensitivity to RAD21 cleavage. To test this, we examined the effect of TSA treatment on CTCF-CTCF loop formation. In DMSO-treated nuclei, we detected strong focal enrichment of interactions between loop anchors, whereas in TSA-treated nuclei, this enrichment remained largely unchanged (Fig. 3g and Extended Data Fig. 8g, left two heatmaps). In salt buffers, TSA treatment slightly weakened loop interactions (Fig. 3g and Extended Data Fig. 8g, third column of heatmaps), and cleaving RAD21 resulted in almost complete loss of loop interactions (Fig. 3g and Extended Data Fig. 8g, second and fourth columns of heatmaps).

To quantify CTCF-CTCF loop strength, we plotted the loop-line for each condition, as defined and calculated by the blue dashed line in the top heatmap of Fig. 1k. Loop-lines showed minimal changes following TSA treatment (Fig. 3h and Extended Data Fig. 8h, left panel). In salt buffers, TSA treatment resulted in slight changes to the loop lines (Fig. 3h and Extended Data Fig. 8h, third panel). Notably, cleaving RAD21 in TSA-treated nuclei caused a significant decrease in loop peaks (Fig. 3h and Extended Data Fig. 8h, second and fourth columns). We further quantified these changes by subtracting the loop lines in low salt buffer without RAD21 cleavage to generate relative loop-lines (Fig. 3i and Extended Data Fig. 8i). The depth of the valley indicates a reduction in loop interaction strength. Cleaving RAD21 in TSA-treated nuclei resulted in a deeper valley (Fig. 3i and Extended Data Fig. 8i, dark solid and dashed lines). Interestingly, we found that all loops, regardless of size, were more sensitive to RAD21 cleavage after TSA treatment (Fig. 4a- c and Extended Data Fig. 9a-c). In summary, these results indicate that in TSA-treated nuclei, cohesin continues to entrap chromatin at CTCF-CTCF loops in the same pseudotopological manner as in DMSO-treated nuclei. Although histone hyperactylation reduces cohesin binding at CTCF sites and increases susceptibility to RAD21 cleavage, the act of encircling chromatin enables cohesin loops to tolerate the chromatin hyperacetylation induced by TSA.

**Fig. 4.**
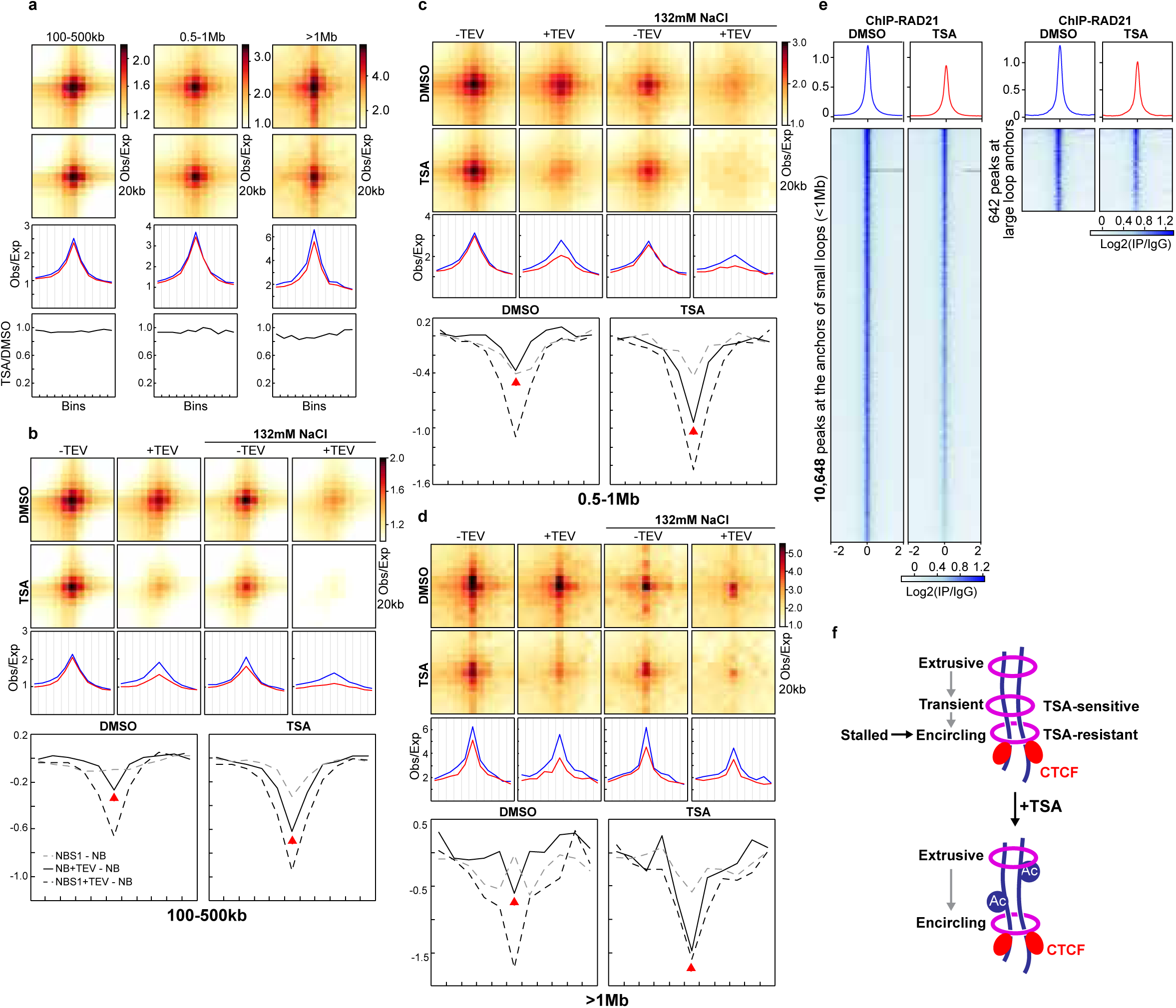
CTCF-CTCF loops tolerate cohesin loss induced by histone hyperacetylation, regardless of loop sizes. a, Averaged Hi-C signals at chromatin loops of three different loop sizes identified in HAP1 cells^12^, and the associated loop-lines and ratios of loop lines as in Fig. 1j**-k**. The Hi-C results are from HAP1 cells treated with DMSO or TSA. **b-d,** Aggregate Hi-C data at loops of different sizes identified in HAP1 cells^12^. Middle and bottom panels indicate loop-lines and difference between loop-lines as in Fig. 3f**-h**. **e**, Average RAD21 ChIP–seq signals in each indicated condition for 642 binding sites that overlapped with the anchors of CTCF-CTCF large loops (1Mb, right) and the rest 10,648 binding sites (left). **f**, Two distinct cohesin populations at CTCF-CTCF sites: a TSA- sensitive, transient form and a TSA-resistant form that encircles pairs of CTCF loop anchors. Histone hyperacetylation selectively dissociates the transitient cohesin. **a-e**, The red and blue lines represent samples treated with and without TSA, respectively.

### Two populations of cohesin at CTCF-CTCF sites

Our analysis of reduced intra-TAD loop interactions indicates that loop extrusion within TADs is disrupted in TSA-treated cells. We initially hypothesized that the longer cohesin extrudes along chromatin, the greater the chance it is lost during TSA treatment, which would preferentially weaken large CTCF–CTCF loops. However, we observed only minimal reduction in interactions at large loops (>1 Mb; Fig. 4a and Extended Data Fig. 7a). One possible explanation is that these loops are held by cohesin that had already stalled at CTCF sites prior to TSA treatment and remained stably bound throughout the 3- hour incubation. Supporting this idea, loops of varying sizes exhibited similar increases in sensitivity to cohesin cleavage, suggesting that stalled cohesin accumulates at CTCF– CTCF loop anchors and encircles pairs of CTCF sites in a stable pseudotopological manner, regardless of loop size (Fig. 4b–d and Extended Data Fig. 7b–d).

We next examined cohesin occupancy at loop anchors of different loop sizes. Large loop anchors showed comparable reductions in cohesin binding relative to small and medium loops, and occupancy of cohesin is also similar (Fig. 4e and Extended Data Fig. 7e). These results indicated that similar amounts of cohesin are lost across loop sizes, while the remaining cohesin continues to associate with chromatin at CTCF sites and maintain looping interactions.

These findings suggest the existence of two distinct populations of cohesin at CTCF– CTCF sites (Fig. 4f). One population appears sensitive to histone hyperacetylation and likely corresponds to cohesin that has not yet encircled chromatin—possibly holding chromatids as an extrusive cohesin within TADs. The other, predominant population encircles the CTCF sites and remains resistant to TSA-induced chromatin hyperacetylation. We refer to the first population of cohesin as transient, TSA-sensitive cohesin, as it transitions toward topologically entrapping chromatin but still exhibits sensitivity to TSA, similar to extrusive cohesin.

This model suggests a two-stage process for cohesin stalling at CTCF–CTCF sites: in the first stage, TSA-sensitive cohesin either arrives at CTCF sites while extruding or is newly loaded there, but remains in a non-encircling state that is susceptible to histone hyperacetylation; in the second stage, chromatids become fully entrapped within the cohesin ring, establishing a more stable, TSA-resistant loop structure.

## Discussion

TSA-induced histone hyperacetylation weakens intra-TAD interactions but leaves CTCF-anchored loops largely unaffected (Fig. 3i), revealing distinct sensitivities between two classes of cohesin-mediated chromatin loops. In our previous work, we showed that cohesin engages chromatin through mechanistically distinct modes: within TADs, cohesin extrudes loops via non-topological engagement, whereas at pairs of convergent CTCF sites, cohesin topologically entraps DNA to form stable, stalled loops^18^. These biophysical differences likely underlie the differential responses to chromatin hyperacetylation.

Extruding cohesin acts through a dynamic cycle of transient chromatin interactions^38–41^. For example, the SMC–RAD21 interface initially clamps chromatin, followed by projection of the SMC1–SMC3 hinge that promotes movement along the chromatin fiber^42^. These transient interactions may be susceptible to changes in chromatin charge and conformation. Histone hyperacetylation alters chromatin surface charge, which appears to result in impared extrusion activity and weakened intra-TAD interactions.

By contrast, cohesin rings that topologically entrap chromatin at looped CTCF–CTCF sites form a more stable configuration, supporting persistent looping interactions that remain largely intact following TSA-induced chromatin hyperacetylation. This resistance highlights the functional and mechanistic distinction between non-topologically and topologically bound cohesin complexes.

Interestingly, despite preserved CTCF-CTCF loops, we observe a marked reduction in cohesin ChIP-seq signal at CTCF loop anchor sites after TSA treatment. This uncoupling between cohesin occupancy and loop interactions suggests the presence of two cohesin populations at loop anchors. We propose that a TSA-sensitive population—likely consisting of newly loaded or transiently extruding, non-encircling cohesin—is preferentially depleted following chromatin hyperacetylation, whereas a TSA-resistant population—composed of topologically bound cohesin—maintains loop interactions. Supporting this model, RAD21 cleavage in TSA-treated nuclei abolishes CTCF-CTCF loops, confirming that maintenance of this type of loops depends on the TSA-resistant, encircling form of cohesin (Fig. 3f-h).

The TSA-sensitive population may reflect cohesin in the process of transitioning between dynamic and stable binding modes. We propose two non-mutually exclusive scenarios. First, at a given loop anchor, CTCF may engage in multiple loop interactions across cells: some stabilized by topologically-bound cohesin holding two CTCF sites, and others formed via non-encircling complexes with distal non-anchor CTCF sites. Consistent with this, we showed in our previous work that only CTCF-CTCF loops are mediated by the topologically-bound form of cohesin, whereas loops where only one anchor is at a CTCF sites are held by the extruding non-topological form^18^. In this scenario, loss of the latter population upon TSA treatment would reduce cohesin ChIP signal, loss of the CTCF- nonCTCF loops, but will not alter aggregate CTCF-CTCF loop strength. Alternatively, paused extruding cohesin may transiently accumulate at CTCF-bound anchors already occupied by stable topologically-bound cohesin. TSA-induced depletion of the extruding population would again reduce cohesin ChIP signal, while preserving the underlying loop that are maintained by the stably bound cohesin complex. Both scenarios account for reduced cohesin occupancy with CTCF-CTCF loop maintenance and highlight the biochemical heterogeneity of cohesin at loop bases.

Together, our findings support a model in which cohesin exists in at least two functionally and biochemically distinct states: a dynamic, TSA-sensitive extruding form that promotes intra-TAD contacts, and a stable, TSA-resistant, topologically engaged form that anchors CTCF–CTCF loops. Histone hyperacetylation selectively disrupts the extruding population while sparing the CTCF-CTCF loop-anchoring form, revealing how chromatin state modulates cohesin engagement and activity.

Our previous results also showed that cohesin at transcription start sites (TSSs) is sensitive to cohesin ring opening, suggesting topological engagement with chromatin at these loci^18^. Given that TSSs are characterized by high levels of histone acetylation, the increased susceptibility of cohesin to RAD21 cleavage likely reflects an open chromatin state that promotes cohesin destabilization. Despite this sensitivity, cohesin at TSSs may play a structural role in stabilizing the transcriptional machinery and facilitating gene regulation. These findings highlight the critical importance of cohesin ring integrity in supporting transcriptional activity at hyperacetylated regions and underscore how chromatin state shapes cohesin dynamics on chromatin.

## Acknowledgements

We thank all the members of both the Liu and Dekker laboratories for discussion. We thank the Deep Sequencing Core at UMass Chan Medical School. We thank the assistance and support from the Biostatistics and Bioinformatics Facility, Genomic Resources Facility, Cell Sorting Facility, and Cell Culture Facility at Fox Chase Cancer Center. Research reported in this publication was supported by the National Cancer Institute of the National Institutes of Health under Award Number P30CA006927. We acknowledge support from the National Institute of General Medical Science (R35GM154879 to Y.L., R35GM144131 to J.R.W.) and W. W. Smith Charitable Trust Grant (C2407 to Y.L.), National Institutes of Health Common Fund 4D Nucleome Program (DK107980, HG011536 to J.D.), the National Human Genome Research Institute (HG003143 to J.D.), the National Cancer Institute (P30CA006927 to J.R.W). J.D. is an investigator of the Howard Hughes Medical Institute.

## Author Contribution Statement

Y.L. and J.D. conceived and designed the project. R.G. and Y.L. performed all the experiments with assistance of H.M.W, K.L.S., C.A. and J.R.W. Y.L. analyzed all the data with assistance of Y.F, R.W. and M.D. Y.L. and J.D. wrote the manuscript.

## Methods

### Cell culture and chemicals

HCT-116-mAID-RAD21 degron2 cells were kindly provided by Yesbolatova, et al, 2020^35^. These cells were cultured in McCoy’s 5A medium, GlutaMAX supplement (Gibco, 36600021) supplemented with 10% FBS (Gibco, 16000044) and 1% penicillin-streptomycin (Gibco, 15140) at 37°C in 5% CO2. HAP1 cells were orginally purchased from Horizon Genomics (Cambridge, UK, C859) and HAP1-RAD21^TEV^ cells were generated as previously reported. Both HAP1 and HAP1-RAD21^TEV^ cells were cultured in IMDM medium, GlutaMAX supplement (Gibco, 31980097) supplemented with 10% FBS (Gibco, 16000044) and 1% penicillin-streptomycin (Gibco, 15140) at 37°C in 5% CO2. Trichostatin A (#T8552) and 5’-ph-IAA (SML3574) were purchased from Sigma-Aldrich, USA and dissolved in DMSO (#D2650) before treating the cells.

### Cell cycle analysis

For cell cycle analysis, cells were washed using 1xDPBS once and fixed in 90% ethanol at -20°C for at least 24 hours. Fixed cells were washed in 1xDPBS and then resuspended in DPBS containing 2mM MgCl2, 0.5mg/ml RNase A (Roche, 10109169001), 1xPI (20xPI in DMSO,). The samples were incubated at 20°C for 30min and then analyzed using an LSRII flow cytometry instrument with green channels to monitor DNA contents. FACS data were processed and analyzed using FlowJo v.3. Viability gates using forward and side scatter were set on each sample. DNA content was plotted as a histogram of the PI channel.

### Nucleus purification and TEV cleavage

Nuclei were purified according to Sanders et al^43^. Briefly, DMSO- or TSA-treated HAP1- RAD21^TEV^ cells were trypsinized, collected in medium, and counted. Around 100million cells were collected and washed in cold PBS twice, and once in NB buffer (10mM PIPES pH 7.4, 2mM MgCl2, 10mM KCl, protease inhibitor (PI, ThermoFisher, #78438)). The cells were then re-suspended in NB buffer with 0.1% NP-40, 1mM DTT, and protease inhibitors, and incubated on ice for 10mins. The cells were lysed using a Dounce homogenizer with pestle A and then loaded onto a sucrose cushion (NB buffer + 30% sucrose + 1mM DTT and 5ml of cell lysis for 20ml sucrose cushion), and finally centrifuged at 800g for 10mins. The nucleus pellets were washed once in cold NB buffer, then resuspended in NB and counted. Around 4 million nuclei were plated onto 60mm poly-D-lysine culture dishes, respectively (Corning BioCoat, #356469), and incubated at 4°C overnight. For the TEV cleavage assay, 60 units of TEV enzyme (AcTEV Protease, Thermo Scientific, 12575-015) were added to 4 million nuclei in 4ml buffer (15U/ml TEV) before the nuclei were plated onto the dishes and kept at 4°C overnight. Control plates contained no TEV, otherwise TEV concentrations were as indicated in Figures.

### Hi-C experiments

Hi-C for fixed cells or the fixed nuclei from the semi-in vitro assay was performed as described in our previous study^18^. Briefly, the fixed cells were first lysed to obtain nuclei. After washed twice with cold NEBuffer 3.1, both the nuclei from fixed cells or from the semi-in vitro assay were 347ul NEBuffer 3.1 with 0.1% SDS and the tube was gently mixed. The tube was then incubated in 65°C for 10min and then put on ice immediately. Before 400U DpnII was added, 40ul of 10% Triton X-100 was added and gently mixed. DpnII digestion was performed at 37°C overnight with gently rocking. Once enzyme digestion was done, the reaction was incubated at 65°C for 15mins to inactivate DpnII. After this, the DNA overhanging ends were filled in with biotin-14-dATP for DpnII digested chromatin at 23 °C for 4 hours and then ligated with T4 DNA ligase at 16 °C for 4 hours. DNA was treated with proteinase K at 65 °C overnight to remove cross-linked proteins. Ligation products were purified, fragmented by sonication to an average size of ∼200 bp and size-selected to fragments of 100–350 bp. We then performed end repair and dA-tailing and selectively purified biotin-tagged DNA using streptavidin beads. Illumina TruSeq adapters were added to form the final Hi-C ligation products, samples were amplified and the PCR primers were removed. Hi-C libraries were then sequenced using PE50 or PE150 bases on an Illumina HiSeq 4000 or an Illumina NovaSeq instrument.

### Hi-C for G1 sorted cells

As previously reported^18^ to sort G1 cells for Hi-C analysis, the cells were fixed following the Hi-C protocol using 1% formaldehyde. The fixed cells were washed in 1×PBS and resuspended in PBS containing 2 mM MgCl2, 0.1% saponin, 0.5 mg ml−1 RNase A and PI (200 μM stock in dimethylsulfoxide). The samples were incubated at 20 °C for 30 min and then analysed using an BD FACS Aria II flow cytometry instrument using the yellow channels to monitor the DNA content. To avoid obtaining any cells in the S phase, only the cells in the left part of G1 peak were collected The FACS data were processed and analysed using FlowJo v.3. Viability gates using forward and side scatter were set on each sample. The sorted G1 cells were then used to generate Hi-C libraries as described as above.

### Hi-C and data analysis

Sequencing data of PE100 or PE150 was first trimmed to PE50 using the in-house scripts. All Hi-C PE50 fastq raw sequencing files were mapped onto hg19 human reference genome using distiller-nf mapping pipeline (https://github.com/mirnylab/distiller-nf). After mapping, aligned reads were further processed to remove duplicates (https://github.com/mirnylab/pairtools) to obtain a set of filtered reads defined as valid pairs. Valid pairs were then binned into contact matrices at 10 kb and 100 kb resolutions using cooler50. Intrinsic Hi-C biases were removed using the Iterative Correctionm and Eigenvector decompositiom (ICE) procedure^44^ was applied to all of the matrices, ignoring the first two diagonals to avoid short-range ligation artefacts at a given resolution, and bins with low coverage were removed using the MADmax filter with default parameters. Contact matrices were stored in ‘.cool’ files and used in downstream analyses.

For aggregation of loop interactions, the previously identified sets of HAP1 looping interactions were used^1,12^. In total, 8334 looping interactions are on the structurally intact chromosomes of HAP1. To visualize the looping interactions, we aggregated 20 kb binned data at all loops using Cooltools. We also aggregated 20kb binned data at different sizes of loops, 100kb-500kb, 500kb-1Mb and greater than 1Mb. The size of a loop refers to the distance between the two loop anchors.

For *P*(s) plots and derivatives, the cis reads from the cooler files were used to calculate the contact frequency *(P)* as a function of genomic separation (s) (cooltools). All of the *P*(*s*) curves were normalized using expected cis interactions (cooltools compute-expected) in each dataset. Corresponding derivative plots were calculated from each *P*(s) plot.

For ratio of TSA-induced P(s) changes in the absence versus the presence of cohesin, the difference of P(s) between DMSO and TSA treatments was first calculated, then the ratio the difference in the absence versus the presence of cohesin was obtained.

To calculate the ratio of TSA-induced P(s) changes in the absence versus the presence of cohesin, the difference in P(s) between DMSO- and TSA-treated samples under each condition was determined. Then the ratio of these differences between cohesin-depleted and cohesin-intact samples was calculated and plotted along genenomic distance.

For interaction aggregation at TAD boundaries, we first calculated observed/expected Hi-C matrices of each sample for 20 kb binned data, correcting for average distance decay as observed in the *P*(s) plots (cooltools compute-expected). We then aggregated the observed/expected Hi-C matrices of each sample at the TAD boundaries that were identified from the sample without any treatments, covering 600kb up and downstream of each boundary, and then generated a pileup heatmap of TAD boundaries for each sample.

For average interactions within TADs, we first identified TAD boundaries by calculating the genome-wide contact insulation score using a 20 kb resolution and boundary calling using the Li threshold (https://github.com/open2c/cooltools). Each TAD region was determined by taking the end position from a TAD boundary and the start position for the subsequent boundary. For each TAD region, the diagonal signal is removed and the number additional bins removed from the diagonal and edges of the TAD are set by the TAD sizes. The intra-TAD intensity is calculated by taking the average contact score of the remaining bins. Outliers in the intra-TAD intensity were removed using the interquartile range. The intra-TAD intensity violin plot was visualized using matplotlib and seaborn.

For compartment analysis, compartment boundaries were identified in cis using eigen vector decomposition on 100 kb binned data with the cooltools call-compartments function. A and B compartment identities were assigned by gene density tracks such that the more gene-dense regions were labelled A compartments, and the PC1 sign was positive. Changes in compartment type therefore occur at locations where the value of PC1 changes sign. Compartment boundaries were defined at these locations, except for when the sign change occurred within 400 kb of another sign change. We noticed that translocation between chr9 and chr22 in HAP1 cells affects compartment assignment on chr9, thus we excluded chr9 for the subsequent compartment analysis for all HAP1 cells.

To measure compartmentalization strength, we calculated observed/expected Hi-C matrices for 100 kb binned data, correcting for average distance decay as observed in the *P*(*s*) plots (cooltools compute-expected). We then arranged observed/expected matrix bins according to their PC1 values of the sample without any treatments in each replicate. We aggregated the ordered matrices for each chromosome within a dataset and then divided the aggregate matrix into 50 bins. Strength of compartmentalization was defined as the ratio of (A–A + B–B)/(A–B + B–A) interactions. Strength of A-A and B-B interactions were separately calculated using AA/AB and BB/BA, respectively. The values used for this ratio were determined by calculating the mean value of the 10 bins in each corner of the saddle plot.

### Western blot for cohesin components

For each condition, 4 millions of nuclei were plated on 60mm poly-Lysine coated plates in 4ml buffer at 4°C for overnight. To analyze cohesin proteins retained in nuclei, all buffers were completely removed from each plate and 200ul RIPA buffer (Thermo Fisher, #89900) containing protease inhibitor and TurboNuclease (Accelagen, #N0103M) were added to each plate. All the plates were then incubated at 4°C for 10minutes and the lysed nuclei were scratched and collected. After spun at 8000g for 10 minutes, the supernatant was transferred to a new tube and 5x sample buffer was added. After mixing, the lysis was boiled at 100°C for 5mins for western blot analysis.

The volume for approximately the same number of cells or nuclei for each sample was loaded into each lane of a protein gel for separation. Two types of protein gel and buffer were used. To separate small proteins (MW<50KD), NuPAGE 4–12% Bis-Tris protein gels (Thermo Fisher, #NP0322BOX) was used with NuPAGE MOPS SDS Running Buffer (Thermo Fisher, #NP0001). For large proteins (MW > 100kD), NuPAGE 3-8% Tris-Acetate protein gels (Thermo Fisher, #EA03752BOX) were used in NuPAGE Tris-Acetate SDS running buffer (Thermo Fisher, #LA0041). Proteins were transferred to nitrocellulose membranes (Bio-Rad, #1620112) at 30 V for 2 h in 1× transfer buffer (Thermo Fisher, #35040) in the cold room. The membranes were blocked with 5% milk in TBST (20mM Tris-HCl, pH 7.4, 150mM NaCl and 0.1% Tween-20) for 30 minutes at room temperature. The membranes were then incubated with the specified primary antibodies diluted 1:1,000 in TBST overnight at 4°C. The membranes were washed three times with TBST for 10 min at room temperature each, then incubated with secondary antibodies (anti-rabbit IgG HRP-linked, Cell Signaling, 7074) diluted 1:5,000 in TBST for 2 hours at room temperature. The membranes were then washed three times with PBS-T for 10 min each. Then, the membranes were developed and imaged using SuperSignal West Dura Extended Duration Substrate (Thermo, #34076) and Bio-Rad ChemiDoc with Image Lab 6.0.1.

### Cell ChIP-seq experiments

ChIP-seq experiments were based on the protocol in our recent work with some minor modifications^18^. Briefly, 30 millions of cells were used for each condition, among which 10 million cells for each antibody. Cells were first washed once using DPBS, then fixed by 10ml DPBS with 1% FA. After rocking for 10min at RT, 540ul of 2.5M glycine were added and continue to rock for 5mins before put on ice for at least 15mins. The cells were then washed once using DPBS before collecting to a 1.7ml tubes from the plates using cell scraper. The cells were spun down at 1000g at 4C for 10min and DPBS was then removed. The pellet was then resuspended in 1ml lysis buffer (20mM Tris-HCl pH 8.0, 85mM KCl, 0.5%IGEPAL, PI) and incubated for 15min. The lysis was then spun at 1000g at 4C for 5min and the lysis buffer was removed. The pellet was then resuspended in 900ul sonication buffer (20mM Tri-HCl pH 8.0, 0.2% SDS, 0.5% sodium deoxycholate, and protease inhibitor). The chromosomes were sonicated to fragments around 200-500bp using QSonica Q800R3 Sonicator (8min sonication time). After spinning at 16,000g for 10min, the fragmented chromatin in the supernatant was split into aliquots of 5 millions nuclei per tube and diluted in 1200ul IP buffer with final 0.1% SDS concentration (20mM Tri-HCl pH 8.0, 150mM NaCl, 2mM EDTA, 0.1% SDS, 1% Triton-100, and protease inhibitor). For each tube, around 4ug of antibody was added and for each antibody, 10 million nuclei were used for ChIP. The primary antibodies used in this study included, CTCF (CST, #3418), RAD21 (Abcam, #ab154769, recognizes N-terminal of RAD21), RAD21 (Abcam, #ab992, recognizes C-terminal of RAD21), Rabbit IgG (Sigma, #I-5006), and WAPL (. Chromatin was incubated with primary antibodies on a rocker at 4°C for overnight. After rewashing with IP buffer, 20ul of Dynabeads Protein G (Thermo Fisher, #10004D) were added to each tube followed by incubation on a rocker at 4°C for 2 hours. After the beads were washed for three times, immunoprecipitated DNA was eluted and 5ng ChIP DNA was used to prepare sequencing libraries in the same way as described for Hi-C above.

### ChIPseq data analysis

Sequencing reads were first trimmed to 50bp long before they were mapped onto hg19 using Bowtie 2. HOMER 4.6 was used to clean mapped reads, examine quality of ChIP experiments, call peaks and generate visualization files^45^. Both profile and cluster plots of ChIPseq signals were generated using Deeptools 3.0.2^46^.

### Boxplot

All the boxplots were drawn using the boxplot() function in R with default settings, the box starts in the first quartile (25%) and ends in the third quartile (75%) with the line in the box indicating the median. The outliers are the data points that are greater than Q3+1.5*IQR (upper outlier), or less than Q1-1.5*IQR (lower outlier). Q1 and Q3 refers to the first and third quartile, respectively. IQR refers to internal-quartile range.

### Statistics and Reproducibility

No statistical method was used to predetermine sample size. Unless specified in the legends, Western blot analyses have been performed at least three times. All Hi-C and ChIPseq analyses have been performed for two biological replicates.

## Data Availability

Deep-sequencing data that support the findings of this study have been deposited in the Gene Expression Omnibus (GEO) under accession codes GSE****. All Hi-C and ChIP-seq libraries are listed in Supplemental Table 2 and 3. Source data have been provided in Source Data. All other data supporting the findings of this study are available from the corresponding author on reasonable request.

## Code Availability

All programs used in this study are from Open Chromosome Collective (Open2C) and are public available in GitHub: https://github.com/open2c. No other customized codes were developed for this study.

---

**Extended Data Fig. 1.**
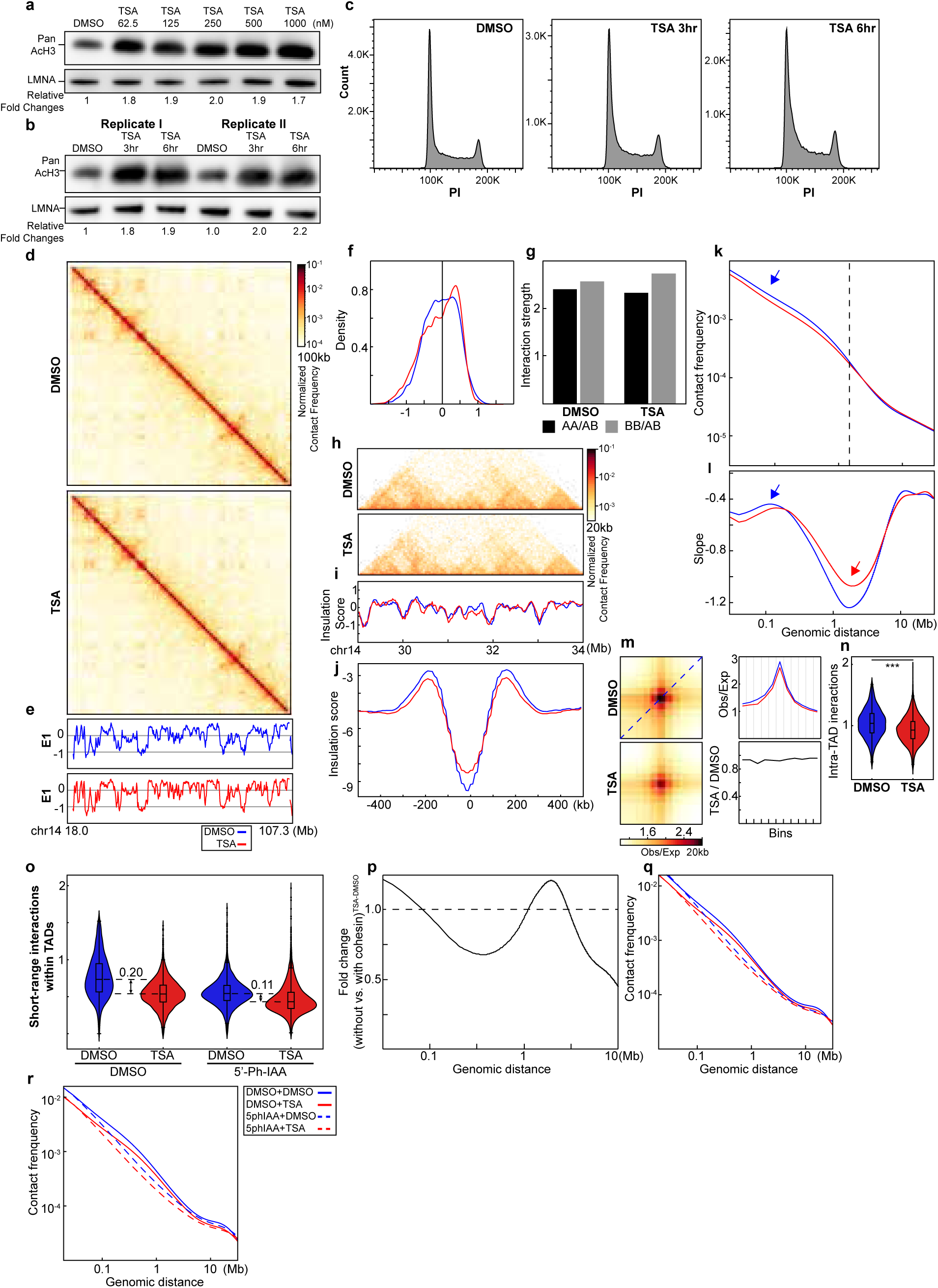
TSA increases levels of histone acetylation and alters chromatin interactions (a replicate of Fig.1) **a**. The levels of acetylated histones by TSA treatments in different concentrations for 3 hours. **b**. The levels of acetylated histones by 500nM TSA treatments for 3 and 6 hours, respectively. Two replicates are shown. **c**. Cell cycle profiles of the cells with TSA treatment for 3 hours or 6 hours. DMSO treatment for 6 hours is included as the control. **d,** Hi-C interaction maps for HAP1 cells treated with/without TSA (top). Data are for the 18.0–107.3 Mb region of chromosome 14. **e**, Eigenvector E1 across the same region as in **a**. **f**, distribution of E1. The red arrow pointed the increased E1 bins after TSA treatment. **g,** Interaction strength of compartments. Dark and grey bars indicate the strength of the A-A and B-B interactions, respectively (see Methods). **h**, Hi-C interaction maps for HAP1 cells treated with/without TSA. Data for the 29–34 Mb region of chromosome 14 are shown. **i**, Insulation profiles for the same region as in **d**. **j**, Aggregate Hi-C data at TAD boundaries identified in the sample treated with DMSO. **k,l**, *P(s)* plots (**k**) and plots of their derivatives (**l**) for Hi-C data from cells treated with/without TSA. The arrow indicates the signature of cohesin loops. **m**, Aggregated Hi-C data at 8,334 loops identified in HAP1 cells according to^12^(upper heatmap). The average Hi-C signals from the bottom-left corner to the top-right corner of the respective loop aggregation heatmaps (top), as illustrated by the blue dashed line in the leftmost Hi-C panel in **m**. This line is defined as the loop-line. The blue and red loop-lines represent loop strength in DMSO and TSA treated samples, respectively. The dark line in the righ bottom plot indicates the ratio between TSA (red) and DMSO (blue) loop-lines. **n**, Average intra-TAD interactions across all TADs (see methods). Blue and red represent DMSO and TSA, respectively. Wilcoxon sum rank test, ***p<0.001. **o**, TSA- induced reduction of cohesin-mediated intra-TAD interactions. **p**, Fold change of TSA- induced *P(s)* alterations in the absence versus the presence of cohesin. **q and r**, *P(s)* plots for two replicates for calculation of fold changes of TSA-induced P(s) alterations in the absence versus the presence of cohesin. The red and blue lines represent samples treated with and without TSA, respectively.

**Extended Data Fig. 2.**
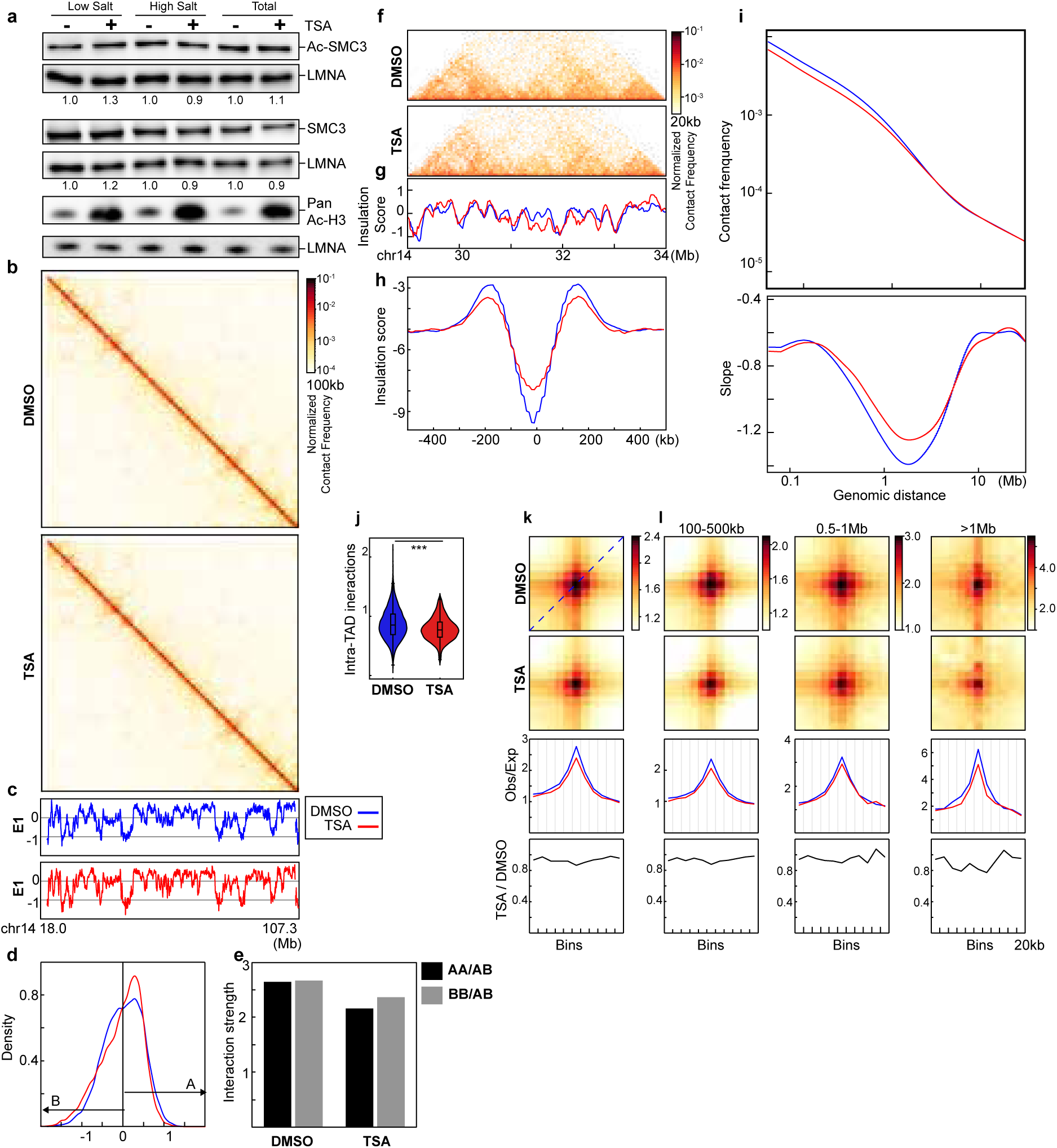
Impact of histone hyperacetylation on wild-type HAP1 cells. **a**, Western blot of SMC3, acetylated-SMC3 and acetylated-Histone 3 in HAP1- RAD21^TEV^ nuclei in low and high salt buffers (100mM and 200mM NaCl). The nuclei were purified from the cells after treated with DMSO or TSA for 3 hours. LMNA was used for normalization. The levels of the SMC3 and acetylated-SMC3 in each condition were normalized to their levels in DMSO-treated nuclei. **b**, Hi-C interaction maps for HAP1 cells treated with/without TSA (top). Data are for the 18.0–107.3 Mb region of chromosome 14. **c**, Eigenvector E1 across the same region as in **b**. **d**, distribution of E1. The red arrow pointed the increased E1 bins after TSA treatment. **e,** Interaction strength of compartments. Dark and grey bars indicate the strength of the A-A and B-B interactions, respectively (see Methods). **f**, Hi-C interaction maps for HAP1 cells treated with/without TSA. Data for the 29–34 Mb region of chromosome 14 are shown. **g**, Insulation profiles for the same region as in **f**. **h**, Aggregate Hi-C data at TAD boundaries identified in the sample treated with DMSO. **i**, *P(s)* plots (top) and plots of their derivatives (bottom) for Hi- C data from cells treated with/without TSA. The arrow indicates the signature of cohesin loops. **j,** Average intra-TAD interactions across all TADs (see methods). Blue and red represent DMSO and TSA, respectively. Wilcoxon sum rank test, ***p<0.0001. **k**, Aggregated Hi-C data at 8,334 loops identified in HAP1 cells according to^12^ (upper heatmap). The average Hi-C signals from the bottom-left corner to the top-right corner of the respective loop aggregation heatmaps (top), as illustrated by the blue dashed line in the leftmost Hi-C panel in **k**. This line is defined as the loop-line. The blue and red loop- lines represent loop strength in DMSO and TSA treated samples, respectively. The dark line in the bottom plot indicates the ratio between TSA (red) and DMSO (blue) loop-lines. **l**, Averaged Hi-C signals at chromatin loops of three different loop sizes, and the associated loop-lines and differential loop lines as in **i**. **c, d, f-m**, The red and blue lines represent samples treated with and without TSA, respectively. Numerical source data and unprocessed blots are provided.

**Extended Data Fig. 3.**
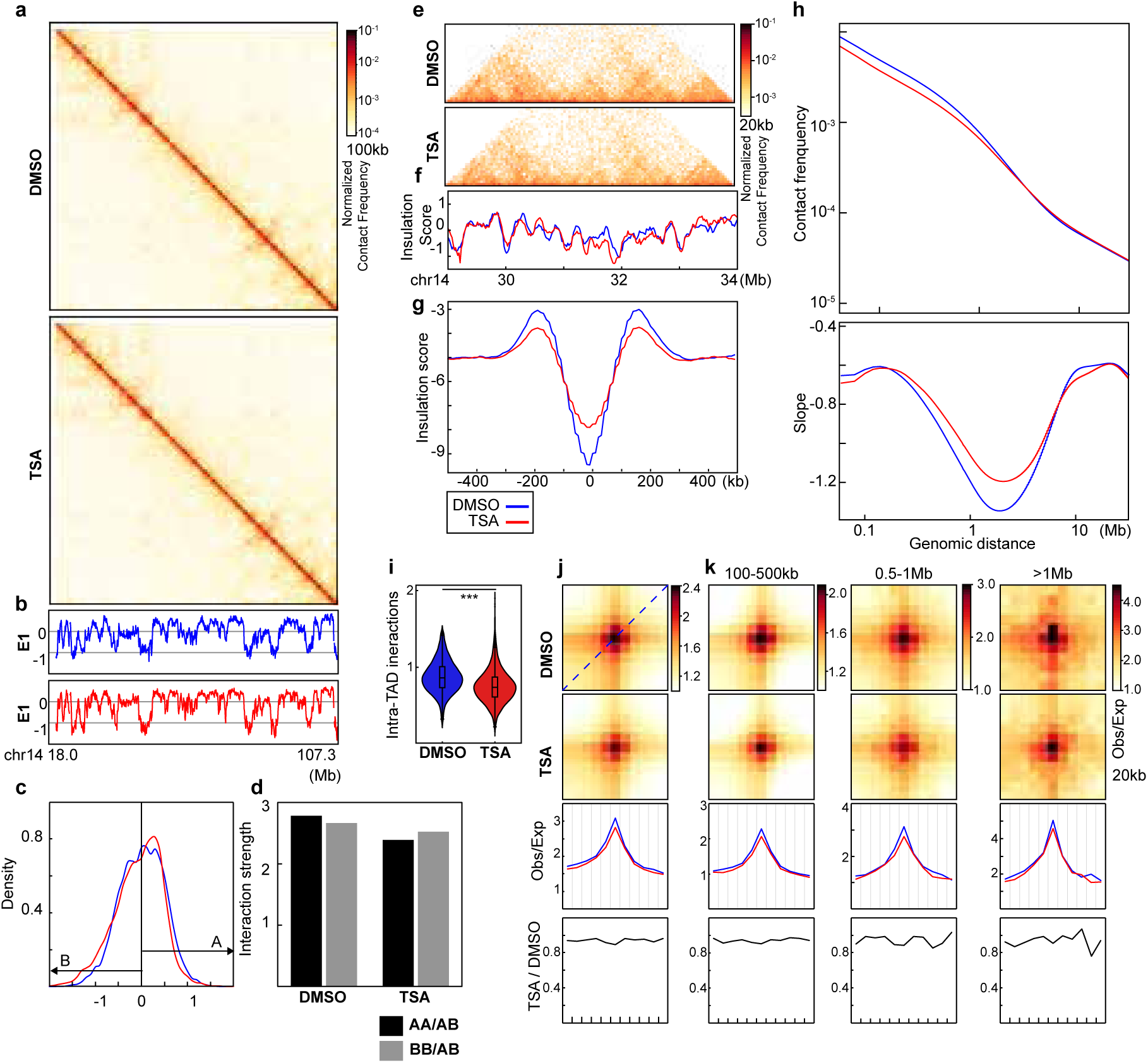
A replicate for impact of histone hyperacetylation on wild- type HAP1 cells. **a**, Hi-C interaction maps for HAP1 cells treated with/without TSA (top). Data are for the 18.0–107.3 Mb region of chromosome 14. **b**, Eigenvector E1 across the same region as in **a**. **c**, distribution of E1. The red arrow pointed the increased E1 bins after TSA treatment. **d,** Interaction strength of compartments. Dark and grey bars indicate the strength of the A-A and B-B interactions, respectively (see Methods). **e**, Hi-C interaction maps for HAP1 cells treated with/without TSA. Data for the 29–34 Mb region of chromosome 14 are shown. **f**, Insulation profiles for the same region as in **e**. **g**, Aggregate Hi-C data at TAD boundaries identified in the sample treated with DMSO. **h**, *P(s)* plots (top) and plots of their derivatives (bottom) for Hi-C data from cells treated with/without TSA. The arrow indicates the signature of cohesin loops. **i,** Average intra- TAD interactions across all TADs (see methods). Blue and red represent DMSO and TSA, respectively. Wilcoxon sum rank test, ***p<0.001. **j**, Aggregated Hi-C data at 8,334 loops identified in HAP1 cells according to^12^ (upper heatmap). The average Hi-C signals from the bottom-left corner to the top-right corner of the respective loop aggregation heatmaps (top), as illustrated by the blue dashed line in the leftmost Hi-C panel in **j**. This line is defined as the loop-line. The blue and red loop-lines represent loop strength in DMSO and TSA treated samples, respectively. The dark line in the bottom plot indicates the ratio between TSA (red) and DMSO (blue) loop-lines. **k**, Averaged Hi-C signals at chromatin loops of three different loop sizes, and the associated loop-lines and differential loop lines as in **j**. **b, c, e-k**, The red and blue lines represent samples treated with and without TSA, respectively. Numerical source data and unprocessed blots are provided.

**Extended Data Fig. 4.**
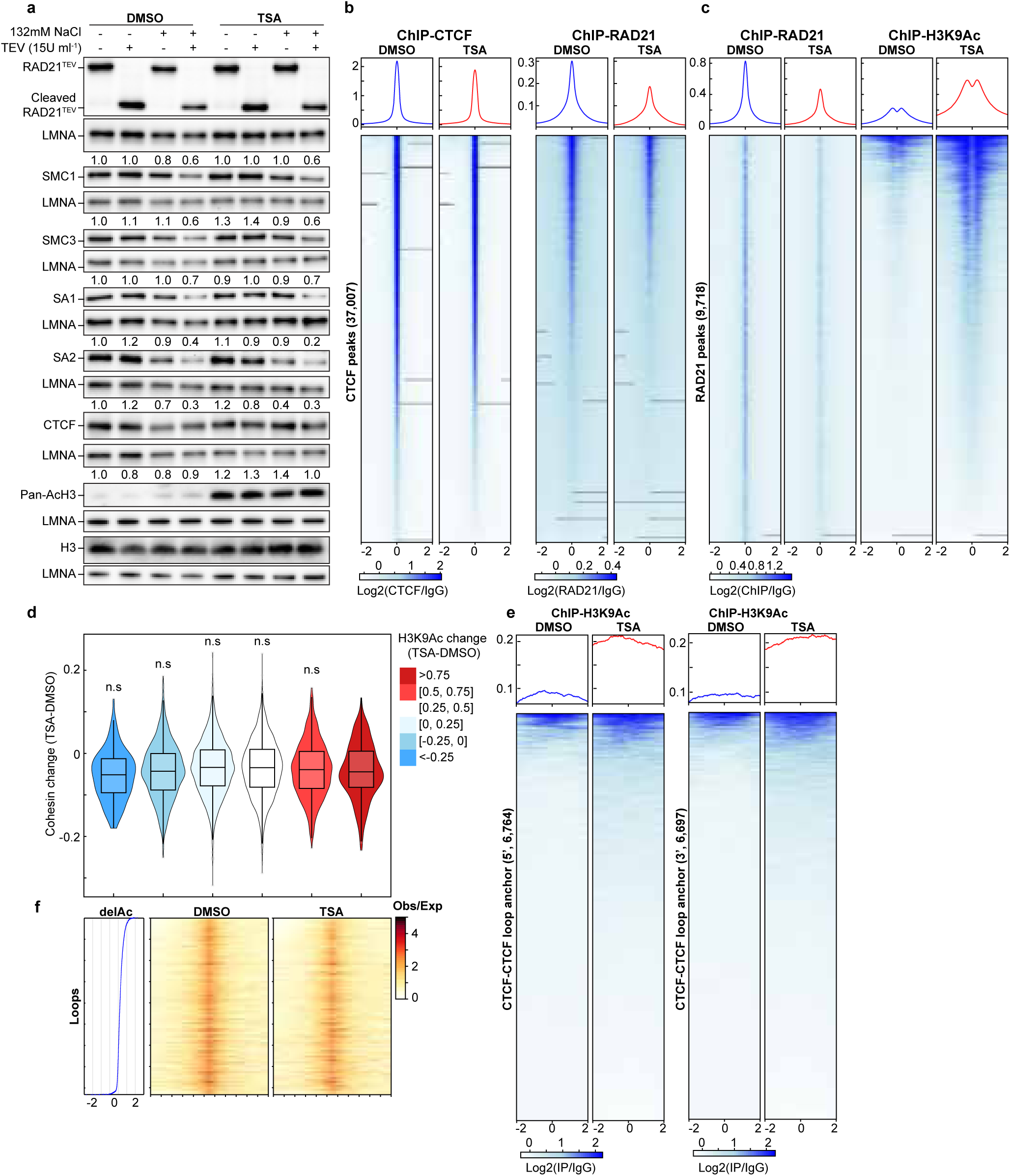
A replicate for impact of histone hyperacetylation on chromatin binding of CTCF and cohesin, and CTCF-CTCF loop sites. **a**, A replicate of Western blot of chromatin retention of cohesin subunits in DMSO- or TSA-treated HAP1-RAD21^TEV^ nuclei in NB or NBS1 buffer (NB buffer + 132 mM NaCl) treated with/without TEV. LMNA was used for normalization (left). The levels of the cohesin subunits in each condition were normalized to their levels in DMOS-treated nuclei in NB buffer without TEV treatment and are indicated in under each LMNA blot. The Western blots in the bottom indicated levels of whole histone H3 and acetylated histones by a pan- acetylated histone antibody. **b**, ChIP–seq signals of CTCF (left) and RAD21 (right) at CTCF binding sites. Average CTCF and RAD21 ChIP–seq signals in each condition for 37,007 CTCF binding sites identified from the CTCF ChIP data from cells treated with DMSO. Heatmaps of CTCF (left) and RAD21 (right) ChIP–seq signals for each condition at each CTCF binding site (bottom). **c**, ChIP–seq signals of RAD (left) and H3K9Ac (right) at RAD211 binding sites. Average RAD21 and H3K9Ac ChIP–seq signals in each condition for 9,718 RAD21 binding sites identified from the RAD21 ChIP data from cells treated with DMSO. Heatmaps of RAD21 (left) and H3K9Ac (right) ChIP–seq signals for each condition at each RAD21 binding site (bottom). **d**, Relationship between RAD21 binding changes and histone acetylation changes. Average of H3K9Ac ChIP-seq signal changes at RAD21 binding sites (+/-2kb) are binned and the changes RAD21 binding strength of each peak in each bin are plotted as a violin plot embed with a boxplot. **e**, ChIP–seq signals of H3K9Ac at 5’- and 3’-loop anchors. There are 6,784 5’- and 6,697 3’- loop anchors, which consist of 8,334 loops. **f**, the relationship between histone acetylation changes and loop line changes. Left ranking blue plot: combined changes of H3KAc ChIP- seq signals at 8,334 pairs of loop anchor sites. Heatmap of ranked 8,334 loop lines for each condition. n.s. not significant, Wilcoxon sum rank test.

**Extended Data Fig. 5.**
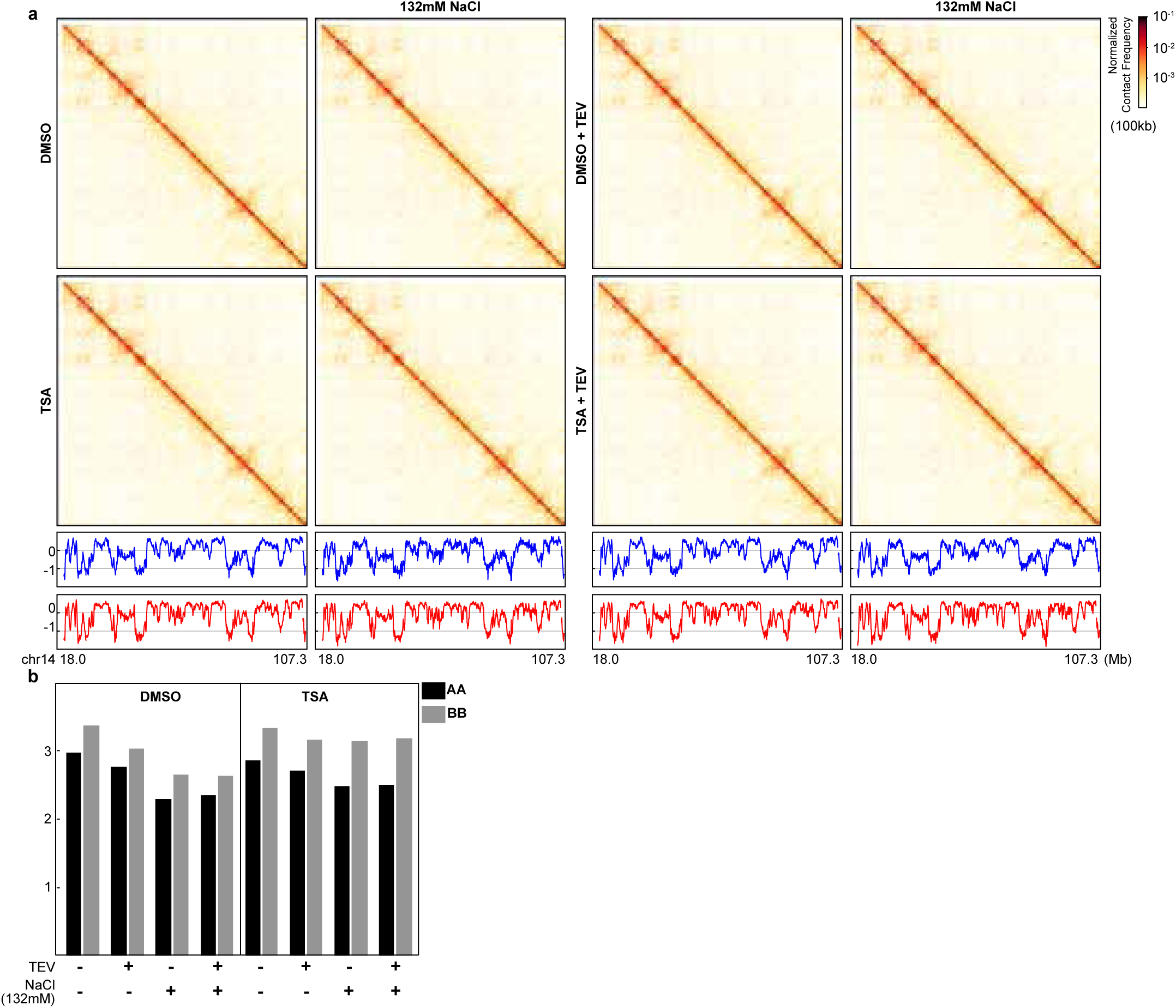
Impact of histone hyperacetylation on compartmentalization of purified nuclei. **a**, Hi-C interaction maps for DMSO- or TAS-treated HAP1-RAD21^TEV^ nuclei treated with/without TEV in NB or NBS buffers. Data are for the 18.0–107.3 Mb region of chromosome 14. Plots with red or blue lines: Eigenvector E1 across the same region as in heatmaps. **b,** Interaction strength of compartments. Dark and grey bars indicate the strength of the A-A and B-B interactions, respectively (see Methods).

**Extended Data Fig. 6.**
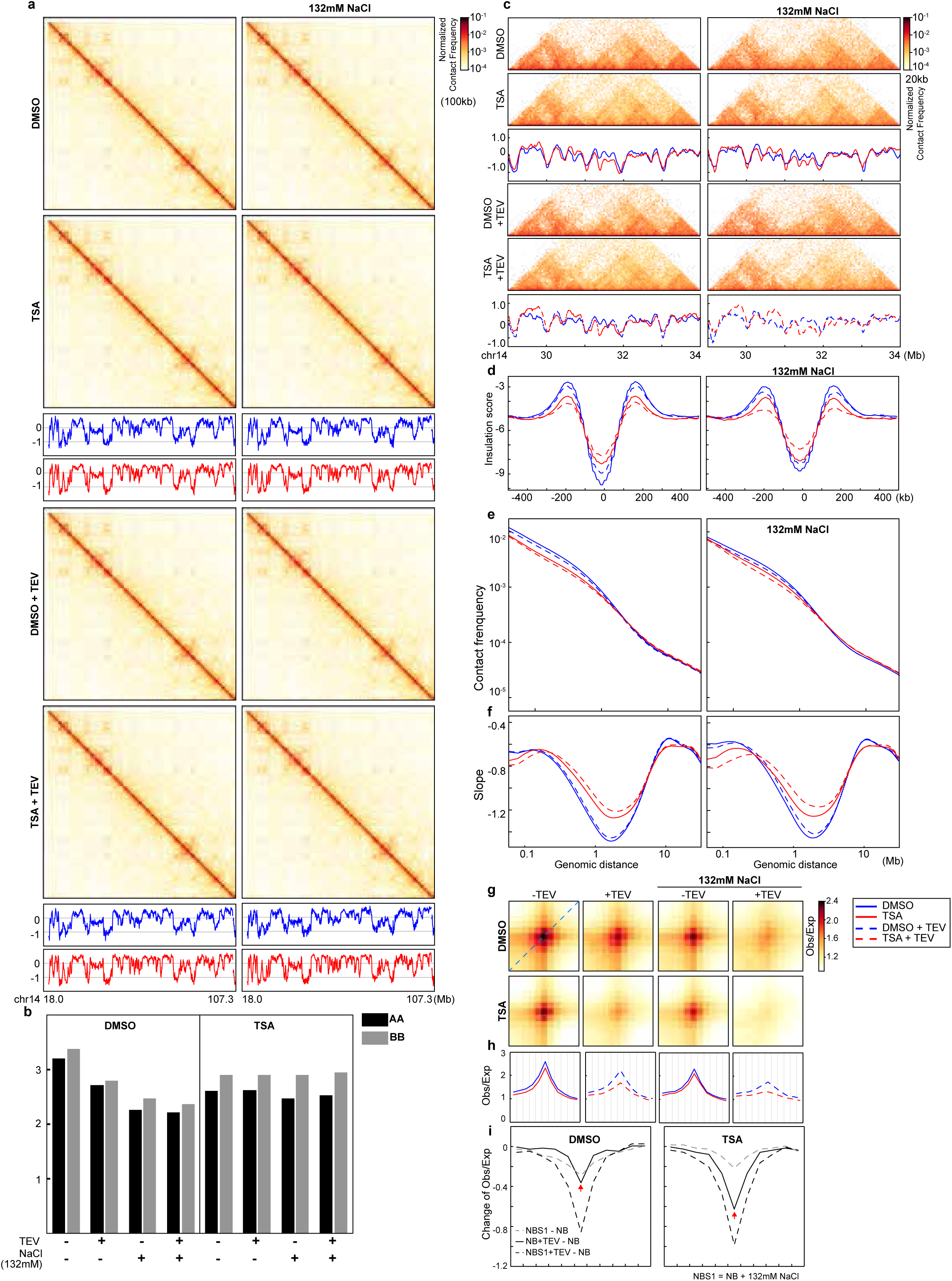
A replicate for histone hyperacetylation increases the sensitivity of cohesin at CTCF-CTCF loop anchors to RAD21 cleavage. **a**, Hi-C interaction maps for DMSO- or TAS-treated HAP1-RAD21^TEV^ nuclei treated with/without TEV in NB or NBS buffers. Data are for the 18.0–107.3 Mb region of chromosome 14. Plots with red or blue lines: Eigenvector E1 across the same region as in heatmaps. **b,** Interaction strength of compartments. Dark and grey bars indicate the strength of the A-A and B-B interactions, respectively (see Methods). **c**, Examples of Hi- C maps obtained with DMSO- or TSA-treated HAP1-RAD21^TEV^ nuclei treated with or without TEV in NB (left) or NBS buffer (right). Insulation profiles (bottom) for the same region for each condition. **d**, Aggregate Hi-C data at TAD boundaries identified in DMSO- or TSA-treated nuclei treated with or without TEV in NB (left) or NBS1 (right) buffer. **e** and **f**, P(s) plots (top) and plots of their derivatives (bottom) for Hi-C data from DMSO- or TSA- treated nuclei with or without TEV treatment in NB (left) or NBS1 (right) buffers. The arrows indicate the signature of cohesin loops. **g**. Aggregate Hi-C data at loops identified in HAP1 cells (as in Fig. 1i; top). The average Hi-C signals from the bottom-left corner to the top- right corner of the respective loop-aggregated heatmaps (top), illustrated by the blue dashed line in the leftmost Hi-C panel in g, are shown (bottom). **h**, The differential loop lines (all other conditions versus NB buffer), left and right panels are DMSO versus TSA treatment.

**Extended Data Fig. 7.**
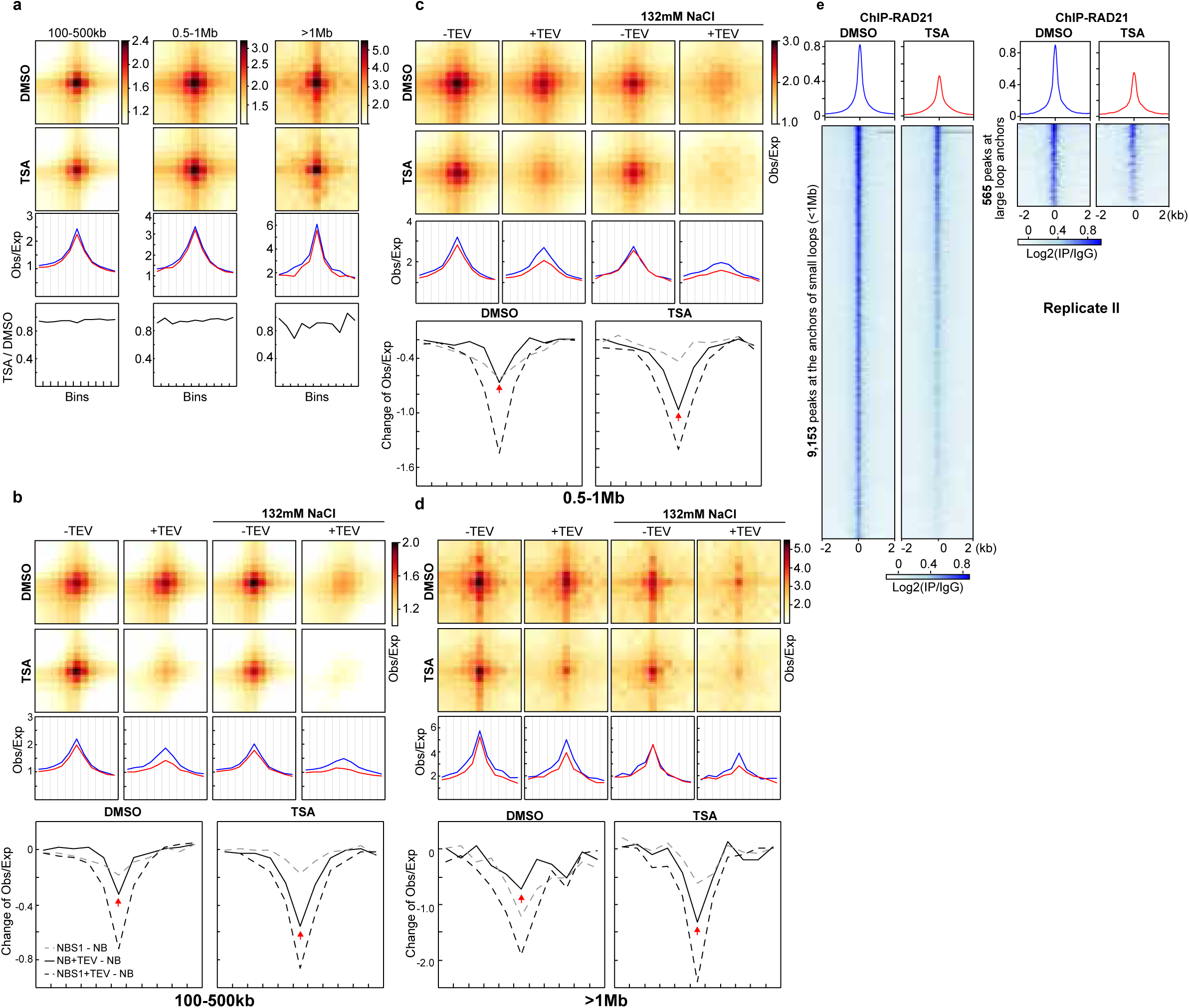
A replicate for CTCF-CTCF loops tolerate cohesin loss induced by histone hyperacetylation, regardless of loop sizes. **a**, A replicate of aggregate Hi-C data at chromatin loops of different sizes identified in HAP1 cells (as in Fig. 4a). For each panel, two upper heatmaps: aggregate Hi-C data at chromatin loops of different sizes; middle: CTCF-CTCF loop lines at chromatin loops of different sizes; bottom: the ratio of loop lines (TSA versus DMSO). **b-d**, A replicate of aggregate Hi-C data at chromatin loops of different sizes identified in HAP1 cells (as in Fig. 4b**-d**). For each panel, upper two heatmpas: aggregate Hi-C data at chromatin loops of different sizes; middle: loop lines at chromatin loops of different sizes; bottom: the differential loop lines (all other conditions versus NB buffer), left and right panels are DMSO versus TSA treatment. **e**, Average RAD21 ChIP–seq signals in each indicated condition for 565 binding sites that overlapped with the anchors of CTCF-CTCF large loops (1Mb, right) and the rest 9,153 binding sites (left). **a-e**, The red and blue lines represent samples treated with and without TSA, respectively.

